# Topological Analysis of Multi-Network Threading in the Pancreas

**DOI:** 10.64898/2026.03.02.708973

**Authors:** Vira Raichenko, Rebin Maaruf, Pia Nyeng, Myfanwy Evans

## Abstract

The pancreas is a complex inner organ that develops through the concurrent formation of multiple interacting biological networks, but the geometric principles governing their spatial organization remain poorly understood. Here, we study pancreatic development by analysing loop structures in ductal, neuronal, and vascular networks using tools from topological data analysis. In particular, we employ the application of chromatic persistence, which is designed to detect topological interactions between distinct subsets of a structure; this is used to quantify how loops from different networks become spatially entangled. Our analysis reveals distinct developmental timelines for loop formation, progressive entanglement between ducts and vasculature, and spatial concentration of threading toward the organ interior, providing a geometric characterisation of coordinated network development in the pancreas.

## Introduction

Developing a deeper understanding of complex biological organs requires methods that can capture both their geometry and the intricate spatial relationships between their constituent systems. Many organs consist of multiple, entangled networks that develop concurrently, interact spatially, and perform distinct but interdependent functions. Quantifying such systems, therefore, calls for frameworks capable of handling complex geometry, topology, and multiscale spatial interactions.

From a biological perspective, we are interested in quantifying organ development through geometric methods. In particular, we focus on the pancreas, whose development and function depend on the coordinated formation of several interacting networks, including ductal, neuronal, and vascular systems. The embryonic ductal system transports enzymes from the pancreas to the intestines, provides the underlying structure of the pancreas, and is the cellular origin of endocrine cells that form the hormone-producing islets of Langerhans (hereafter named islets). The pancreatic neuronal system is required for the regulation of hormone secretion from islets, while the vascular system provides nutrients and allows pancreatic islets to regulate blood glucose levels. A deeper understanding of the coordination of these three systems is thus motivated by the need to better understand pancreatic diseases such as diabetes, where these functions are affected.

The pancreatic duct is a branched tube that forms through a series of epithelial transformations, from a bud, to a plexus, which eventually gives rise to the ductal tree [1]. Pancreatic endocrine differentiation is tightly coupled to the growth and morphogenesis of the developing duct [2]. We have recently demonstrated that ductal loops that appear during the plexus stage, form unique niches for endocrine differentiation, and that they are removed by a novel loop closing mechanism, emphasising that understanding duct-architecture is essential for explaining the emergence of pancreatic function [1]. Pancreatic innervation begins with the formation of an intrinsic neuronal network by invading neural crest cells. The intrinsic neural network forms connections with islets, vasculature, and extrinsic neuronal fibers from the central nervous system [3]. Interactions between neurons and the epithelium may also shape duct morphogenesis and islet differentiation [3]. The anatomy and functional roles of the mature pancreatic innervation, including sympathetic, parasympathetic, and sensory pathways, have been characterized recently by 3D imaging and linked to metabolic disease [4]. The vascular network plays a similarly critical role in development and function, providing not only nutrition, but also signals that regulate pancreas ductal branching and endocrine differentiation [5]. Vasculature in the pancreas have been studied previously, including detailed analyses of vascular organisation [6] and 3D reconstructions of pancreatic vasculature [7]. However, these studies do not address vascular development longitudinally across development, nor do they investigate topological features of the vascular network, or coordination with neuronal and ductal systems. In conclusion, while individual pancreatic networks have been examined in isolation, their joint spatial organisation during development has to our knowledge not yet been addressed. Of particular interest, a whole-organ 3D geometric atlas of adult human and mouse pancreatic islets and their spatial relationships to neural networks demonstrates that innervation is not uniformly distributed but instead biased toward large endocrine structures and reorganised in disease [4]. Their work highlights that pancreatic function is encoded in the geometry and relative placement of distinct structural systems rather than in isolated cell types. Our work builds on this perspective by explicitly focusing on loop formation and topological interactions between neuronal, ductal, and vascular networks over developmental time.

In this study, we analyse spatial interactions between networks with different biological functions. Such interactions are inherently geometric and topological, as they involve proximity, entanglement, and the formation of loops and cycles across systems, Figure 1. These questions naturally call for tools from topological data analysis (TDA), which are designed to capture robust topological features of complex data.

**Fig 1.**
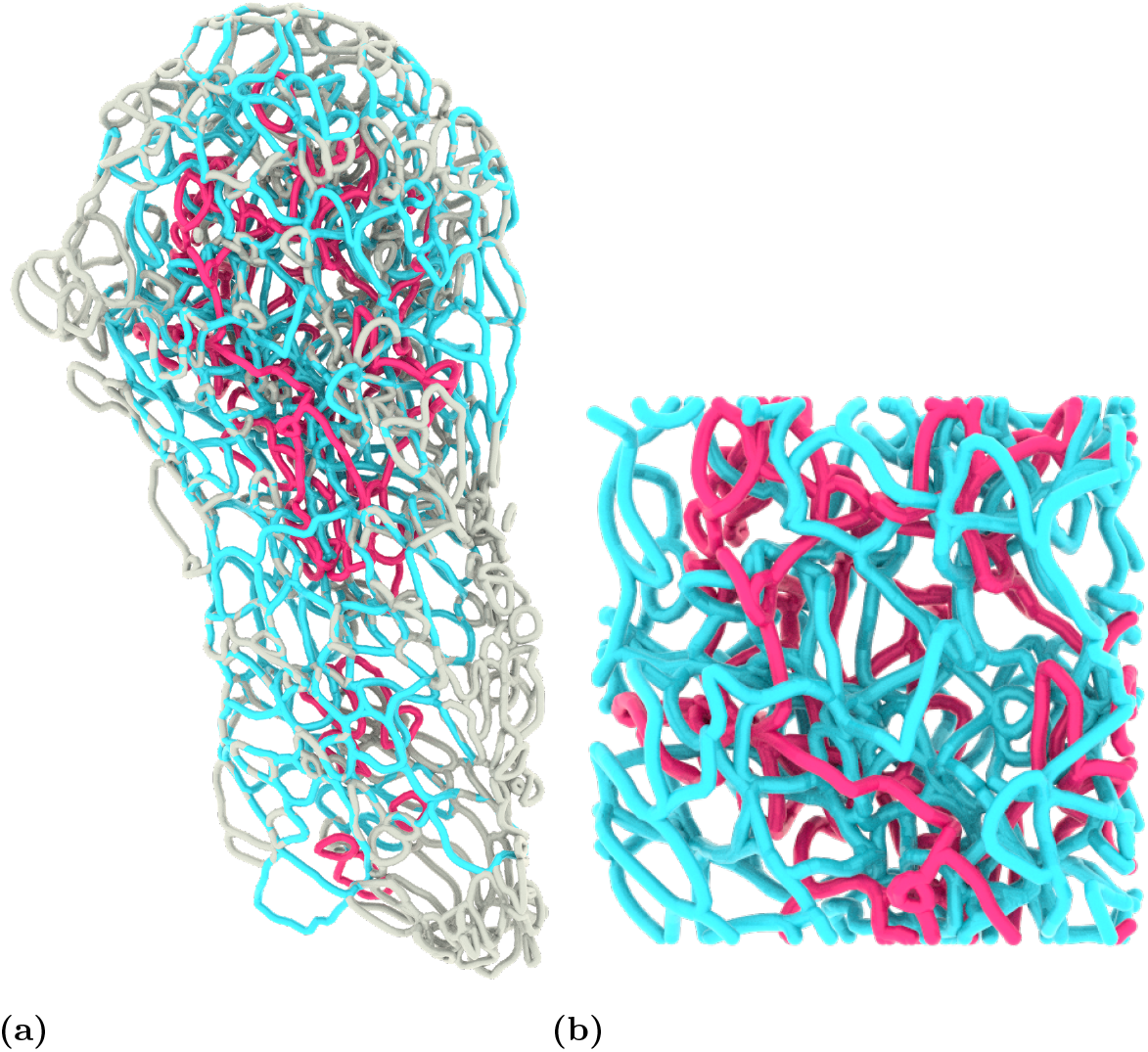
Loops from DUC (ducts) and VAS (vasculature) entangled together. (a) Interacting (entangled) regions of the network, composed of loop structures, are highlighted in pink (DUC) and blue (VAS). White loops correspond to regions where loops from both networks are present but do not exhibit direct interaction. (b) A close-up view of one entangled region, illustrating the spatial interweaving and local geometric relationships between loops from the two networks.

Persistent homology [8–10] provides a mathematically rigorous characterisation of the shape and topology of a structure across scales. Its stability with respect to noise and deformations makes it particularly well-suited for biological data, where measurements are often imperfect and structures exhibit significant variability. When paired with statistical analysis, persistent homology has become a central tool in TDA. At the same time, persistent homology enables the identification and characterisation of concrete geometric features associated with topological changes; in this work we focus in particular on the extraction of loop formation events, as we have previously highlighted the importance of ductal loops [1].

When studying the spatial relationships between multiple interacting structures, classical persistent homology is not sufficient, as it does not distinguish between features arising from different components. This has motivated the development of kernel and image persistence [11], also referred to as chromatic TDA in more recent work in the setting of alpha complexes [12]. Chromatic persistence enables the analysis of topological features while preserving information about the origin or type of each component, making it particularly suitable for studying interactions between multiple biological networks, Figure 1.

The robustness and interpretability of persistent homology have made it an effective tool for extracting scientific insight from complex data across diverse domains. In geoscience, topological descriptors of pore structure in sandstones predict capillary fluid trapping more reliably than classical network statistics [13]. Applied to soft biological materials, persistent homology reveals microstructural gradients in the fibrous architecture of silkworm cocoons that conventional imaging analysis cannot resolve [14]. Persistence barcodes serve as structural fingerprints for classifying nanoporous materials with distinct pore geometries [15], while in zeolitic imidazolate frameworks, the accumulative persistence function links medium-range atomic order to thermal stability [16]. Beyond porous media, persistence diagrams of cell contours detect morphological sub-populations in heterogeneous stem cell cultures, outperforming classical shape descriptors [17].

Topological approaches to spatial interaction have already shown promise in a variety of applications, including tumor analysis [18]. In the context of vascular systems, persistent homology has been used to quantify vessel loops and their spatiotemporal distribution, providing multiscale descriptors that surpass standard single-scale morphological measures [19]. In biology, such questions are often driven by the need to understand how different cell types or structures influence each other through their spatial organization, for example in the tumor immune microenvironment, where spatial configuration has been shown to impact disease progression and treatment response [20]. Related ideas also appear in studies of entangled ring structures [21], highlighting the relevance of topological descriptors for complex intertwined systems.

In this paper, we extract loop structures from segmented biological 3D image data using TDA-based methods. We then apply chromatic persistence to quantify loop formation and spatial relationships in the developing mouse pancreas. Our analysis focuses on the topology and geometry of three interacting networks: neurons, ducts, and vasculature. Throughout this work, we denote these networks as NEU (neurons, as detected with a anti-GAP43 antibody), DUC (ducts, as detected with an anti-MUC1 antibody), and VAS (vasculature, as detected with an anti-PECAM antibody). The analysis summarises our progress on characterising loop formation within each network and on quantifying their spatial interactions during pancreatic development, encompassing embryonic ages E12.5, E13.5 and E14.5 in the mouse. We observe that loop formation does not occur simultaneously across all networks: loops in the ductal and vascular networks are already present at E12.5, whereas neuronal loops mostly appear at E14.5 with just a few formed at E13.5, suggesting that the neuronal network establishes its loop architecture after the ductal and vascular scaffolds are in place. The vascular network maintains a stable relative loop density throughout development, indicating that it does not remodel toward a tree-like topology, while the ductal network shows a decrease in loop density from E13.5 to E14.5, consistent with remodeling toward a more tree-like structure. Characteristic loop sizes are similar across all three networks, ranging from approximately 63 to 81 *µ*m, and their size distributions stabilise by E13.5. Loops in both ducts and vasculature are spatially concentrated toward the centre of the organ, with no correlation between loop size and position. Applying chromatic persistence to quantify inter-network entanglement, we find that the neural network is highly entangled with vasculature, with approximately 84% of neuronal loops threaded by blood vessels at E14.5 compared to 58 % by ducts, suggesting that the vascular network provides the dominant spatial scaffold for neuronal development. Ductal-vascular entanglement is mutual and progressive, with approximately 40–50% of loops in each network threaded by the other at E14.5, increasing steadily from E12.5. In contrast, the neuronal network contributes minimally to threading of other networks, with only 5% of ductal and 15% of vascular loops threaded by neurons. Across all network pairs, larger loops are preferentially threaded, and threading is concentrated in the organ interior, where loops closer to the centre are more frequently entangled while those near the periphery tend to remain non-threaded.

## Materials and methods

### Animal Studies

Mice were housed at the Department for Experimental Medicine at University of Copenhagen. All experiments and breeding was approved by the Animal Experiments Inspectorate (Ministry of Food, Agriculture and Fisheries of Denmark). For each embryonic stage, a male C57BL/6BomTac mouse was set up for timed mating with 2 female C57BL/6BomTac mice in the afternoon, and the presence of a vaginal plug was checked every morning for up to 2 days. The day of vaginal plug was recorded as E0.5. Embryos were collected at day E12.5 (n=5), E13.5 (n=3) and E14.5 (n=3), at which time anatomical features such as limb morphology and crown-rump length were used to verify the developmental stage. Stomach, duodenum, spleen and pancreas were dissected in one piece from embryos at noon on the indicated day and immediately fixed in 4% paraformaldehyde at 20 °C for 2 to 16 hours depending on the size of the tissue and transferred through an ascending gradient to 100% methanol for storage at −20 °C. For a more detailed description of embryonic mouse pancreas dissection and processing, see (Heilmann et al., 2021) [22].

### Tissue preparation and whole mount immunofluorescent staining

Throughout the procedure samples were kept intact in 2 mL round-bottom Eppendorf tubes. Samples were first bleached in a 1:1:4 dilution of 30% H2O2:DMSO:MeOH (Dent’s Bleach) for 2 hours at 20 °C. They were then rehydrated through a descending gradient of methanol to 1x phosphate buffered saline with 0.01% Triton-X100 (PBST), and blocked for 2 hours at room temperature in a TNB blocking buffer (Perkin Elmer). Primary antibodies for GAP43 (rabbit anti-GAP43, dilution 1:500, Abcam Ab16053), MUC1 (Armenian hamster a-MUC1, dilution 1:500, Thermo Scientific MA-11202), and PECAM (Goat anti-CD31/PECAM, dilution 1:500, Bio-Techne AF3628-SP), were diluted in blocking buffer as indicated and added to the samples, which were incubated at 4 °C for three days on a rocking bed. Thereafter, samples were washed 4 times in 1x PBST for one hour each. Secondary antibodies Donkey Alexa Fluor 488-anti-Rabbit FAB fragment AB2340620 and Donkey Rhodamine REd-X-anti-goat whole IgG AB2340423 from Jackson Immunoresearch were diluted 1:1000 in blocking buffer and added to samples, which were incubated overnight at 4 °C on a rocking bed. Samples were then washed three times in PBST for 20 minutes each. Secondary antibody Goat Alexa Fluor 647-anti-Armenian Hamster whole IgG AB2339001 from Jackson Immunoresearch was diluted 1:1000 in blocking buffer and added to samples separately (so as not to react with unbound anti-Goat secondary). Samples were incubated overnight at 4 °C on a rocking bed. Samples were then washed three times in PBST for 20 minutes each. Finally, samples were dehydrated through an ascending gradient to 100% methanol, and cleared and mounted in 1:2 benzyl alcohol and benzyl benzoate (BABB) in a cavity glass slide with a #1.5 cover glass for imaging.

### Image acquisition

A Zeiss LSM780 laser scanning inverted confocal microscope equipped with a Plan-Apochromat 25x/0.8 NA 25X multi-immersion objective and GaAsP detector was used for 3D imaging at room temperature. Images of the dorsal lobe of the pancreas were acquired as tiled Z stacks of 1024 × 1024 pixel and 180-300 z grey-scale images with 10% overlap between tiles. Each voxel is 0.332 × 0.332 × 1.4*µm* (x,y,z).

### Image processing and segmentation

Tiled images were stitched using Zen Black software and exported as an lsm file. The lsm file was converted to an ims file using Imaris software and segmented in Imaris: Each dataset was cropped in three dimensions to remove non-pancreatic tissue, and each channel was segmented using the Surfaces function with identical threshold settings for all samples. Following segmentation, each surface was used to create a binary image by masking. The binary image Z-stack with three channels was exported as a TIFF stack using Open Microscopy Environment TIFF format. As a result, we ended up with segmented masks of three networks: neurons labeled with antibody for the protein GAP43(**NEU**), ducts labelled with an antibody for the protein MUC1(**DUC**), and vasculature labeled with an antibody for the protein PECAM(**VAS**). Each net was processed in the same way with the following steps: cleaning, skeletonisation, and loop extraction.

### Cleaning and Skeletonisation

For skeletonisation, we use a combination of discrete Morse skeletonisation computed using the diamorse library [23, 24] and MATLAB’s `bwskel` function, which implements the medial-axis transform [25]. When the images are too large, we perform the computation on two separate subvolumes split along the y-axis, with an overlap of 300 voxels. When merging the results, we take the union of the Morse skeletons in the overlapping region to ensure that no connections are missed.

The Morse skeleton provides a topologically correct thin structure, but some regions remain locally thickened. Here, we use simplification with the persistence limit for feature cancellation (-p) set to 10 to make the Morse skeleton as thin as possible. By applying the medial-axis algorithm, the remaining thick regions are resolved, resulting in a thin skeleton with preserved loop structure.

The voxelised skeleton is then converted into a graph by connecting voxels using 26-neighbourhood adjacency, where the neighbourhood of a voxel consists of the 26 surrounding voxels that share a face, edge, or vertex with it. Loops are extracted by computing a minimum cycle basis [26] using the NetworkX Python package [27]. This procedure produces a large number of loops, many of which are artefacts of the Morse skeleton construction (see Figure 2).

**Fig 2.**
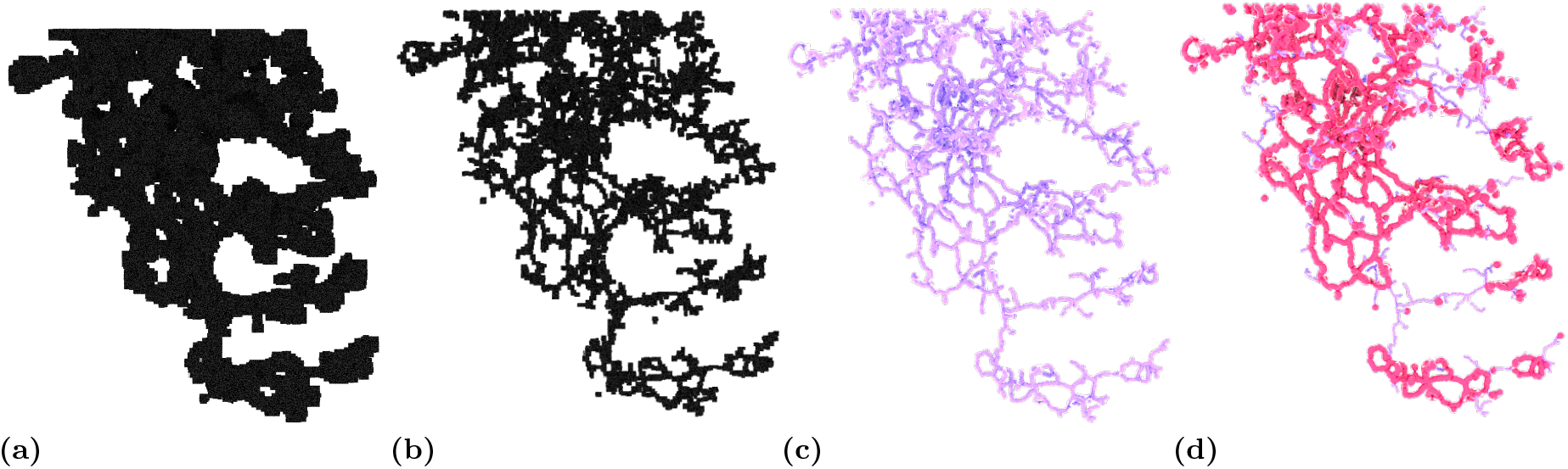
Displaying the processing steps from raw voxelized data to loop extraction. The example is shown on a piece of DUC (ducts). (a) 2D rendering of the raw voxelized data (note that loops in the z plane will not be visible in this rendering). (b) Morse skeleton. (c) Skeleton. (d) All loops extracted from the skeleton in (c), shown in pink.

### Loop extraction

To extract the loops from the data, we use Persistent Homology. We set the networks to be black and the background to be white, and we compute Signed Euclidean Distance Transform(SEDT) with the corresponding spacing that accounts for the physical dimensions of the image, i.e. each voxel is assigned the signed distance to the interface. The SEDT serves as a filtration function that begins inside the body of each network and grows outward, so that the initial level sets capture the geometry of loop formation within the network itself. To compute the persistent homology that records all topological changes throughout this filtration, we use the diamorse library. Those points in the persistence diagram of dimension 1 that have negative birth and positive death represent *true* loops from the data, see Figure 3.

**Fig 3.**
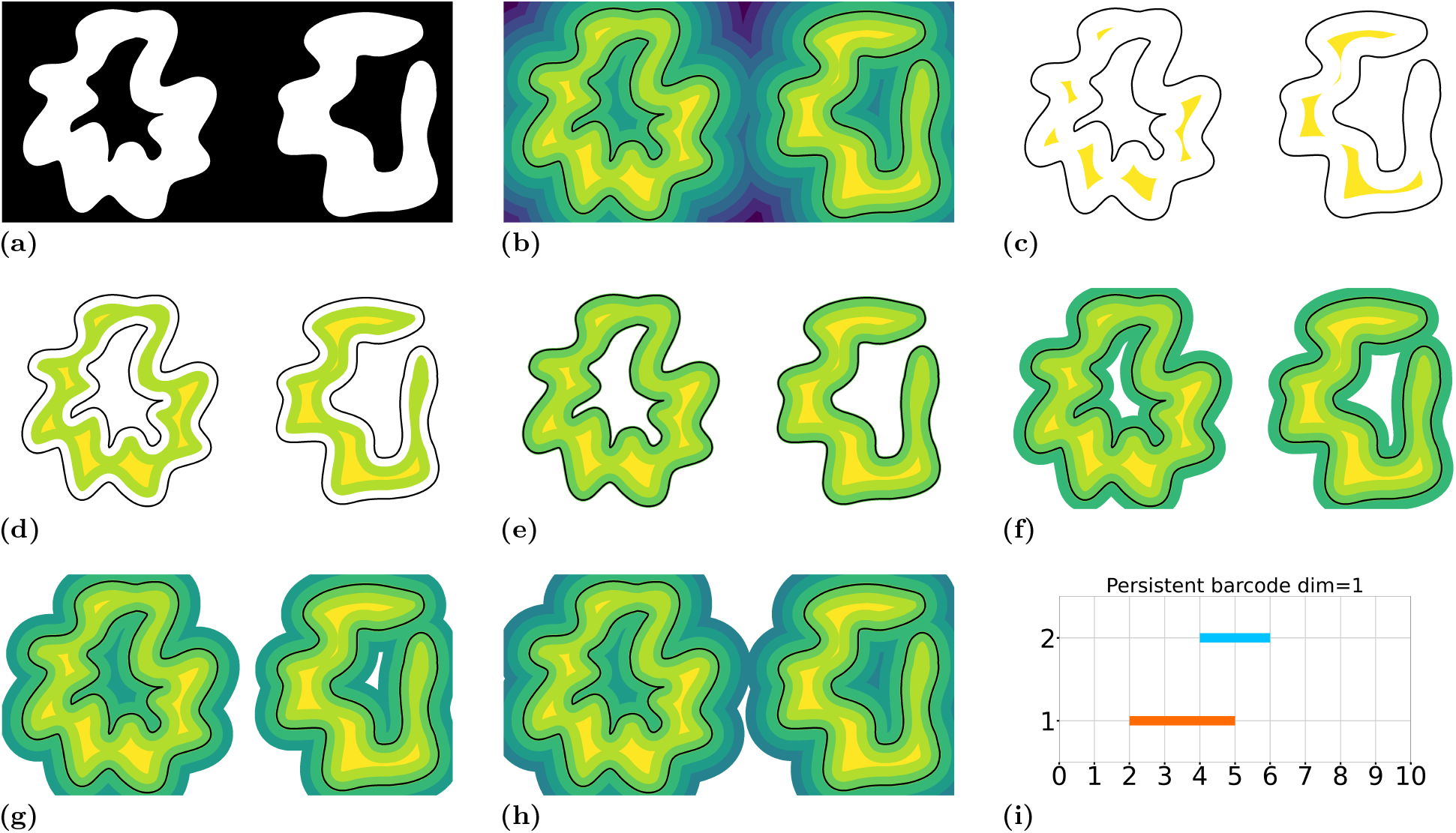
A 2D illustration of a persistence barcode on loop-like data. (a) The input mock image. In (b)–(h), the black outline shows the boundary of the white set in (a). The distance map takes positive values inside the black outline and negative values outside it. (b) (b) SEDT filtration of image (a). All white pixels from (a) are assigned a positive value equal to their distance to the boundary (black outline), while all black pixels are assigned a negative value of their distance to the boundary. The resulting Signed Euclidean Distance Transform is then visualised as a binned level set, where each color band represents a range of distance values. (c)–(h) A sequence of nested subsets of the filtration. We show the first six steps in which non-trivial topological changes occur. Each step contains the set from the previous step. In (d), the first loop is formed; in (f), the second loop is formed; in (g), the first loop becomes filled and no longer exists; and in (h), the second loop becomes filled. (i) The persistence barcode, showing in two dimensions the topological events that occur in (c)–(h). The orange bar corresponds to the first loop on the left—the “true” loop. The blue bar corresponds to the second loop on the right, which does not have loop-like topology in the underlying data.

Using the information from the persistence diagram, we select loops captured by persistent homology as “true” loops in the data. For our dataset, we select points from the 1-dimensional persistence diagram whose birth value is negative (*b <* 0) and whose death value exceeds 2 (*d >* 2). The condition *b <* 0 and *d >* 0 identifies points representing true loops; however, we set *d >* 2 to exclude small loops arising from imaging artefacts rather than genuine structural features. We pair these persistence points with geometric loop representatives extracted from the skeleton. Each persistence point is associated with a birth voxel, which lies on the loop boundary, and a death voxel, which lies inside the loop. We use this information to construct a pairing procedure as follows.

- **Candidate harvesting**. The union of these two following sets forms the candidate pool. For a given persistence point, we first harvest candidate loops from two sources:
  i. loops whose voxels lie within a neighbourhood of the birth point, using *k*-nearest neighbour search with *k* = 768, augmented by radius expansion up to 128 voxels if fewer than 20 candidates are found;
  ii. loops intersecting a sphere centred at the death point with radius |*d* |+ 25, where *d* is the death scalar (the SEDT value at the death voxel).
- **Hard geometric constraint**. All candidates must satisfy the constraint that the minimum distance from any loop voxel to the death point does not exceed |*d* |+ 25. If no candidate satisfies this constraint, it is relaxed and the closest candidate is reported.
- **Spherical arc length**. For each candidate loop, we project its vertices onto the unit sphere centred at the death point and compute the total geodesic arc length Ω of the resulting spherical curve. This quantity measures the spherical arc length of the loop as seen from the death point: a loop that wraps closely around the death point projects onto a long curve on the sphere, producing a large Ω, whereas a distant loop that does not enclose the death point produces a short projected curve. Since Ω measures perimeter rather than enclosed area, it is well-defined for non-convex loops and self-intersecting projections, and can naturally exceed 2*π* (the case for a perfect circle around the centre of the sphere).
- **Selection rule**. We define a spherical arc length threshold Ω_0_ = 5.0 radians. If any candidate has Ω *≥* Ω_0_, we select among those candidates the one with the smallest centre distance (distance from the loop’s barycentre to the death point). If no candidate reaches this threshold, we fall back to selecting the candidate with the largest spherical arc length, with ties broken by smallest centre distance. **Remark.** This procedure does not guarantee a perfect pairing; certain loop geometries can lead to mismatches. Since the loops are embedded in three dimensions, establishing ground-truth correspondences by direct inspection is impractical, and no independent validation set is available. Nevertheless, visual comparison of the paired loops with their assigned persistence points shows consistent spatial agreement across the dataset (see Figure 4), suggesting that the method produces reliable pairings in practice
- **Injectivity and collision resolution**. The initial pairing is non-injective: multiple persistence points may select the same loop. We resolve such collisions through three sequential stages. Stage 1: Label collision resolution. When multiple births select the same loop, we attempt to reassign some births to alternative loops. A birth may move to an alternative only if: (i) the alternative has Ω *≥* 5.0, and (ii) the increase in centre distance satisfies Δ_center_ *≤* 18.0. Among colliding births, one “keeper” remains on the contested loop—chosen as the birth that either has no acceptable alternative or would incur the largest centre-distance penalty if moved. This process runs for up to 2 passes. Stage 2: Same-death resolution. When multiple births share the same death voxel (identified by rounding coordinates to 3 decimal places), we resolve collisions deterministically. The birth whose birth point is closest to the loop’s voxel set is designated the keeper. Other births are moved to their next-best alternative, provided that alternative has Ω *≥* 5.0; otherwise, the birth is marked as unpaired. Stage 3: Strict final cleanup. Any remaining collisions are resolved by retaining exactly one birth per loop and marking all others as unpaired. The winner is selected using the same decision logic: if the birth has any candidate with Ω *≥* 5.0, prefer smallest centre distance; otherwise, prefer largest spherical arc length. After these stages, the pairing is injective, though some persistence points may remain unpaired. The example of final matching is shown in Figure 4. A demonstration of pairing on a small example is provided in the Supporting Information.

**Fig 4.**
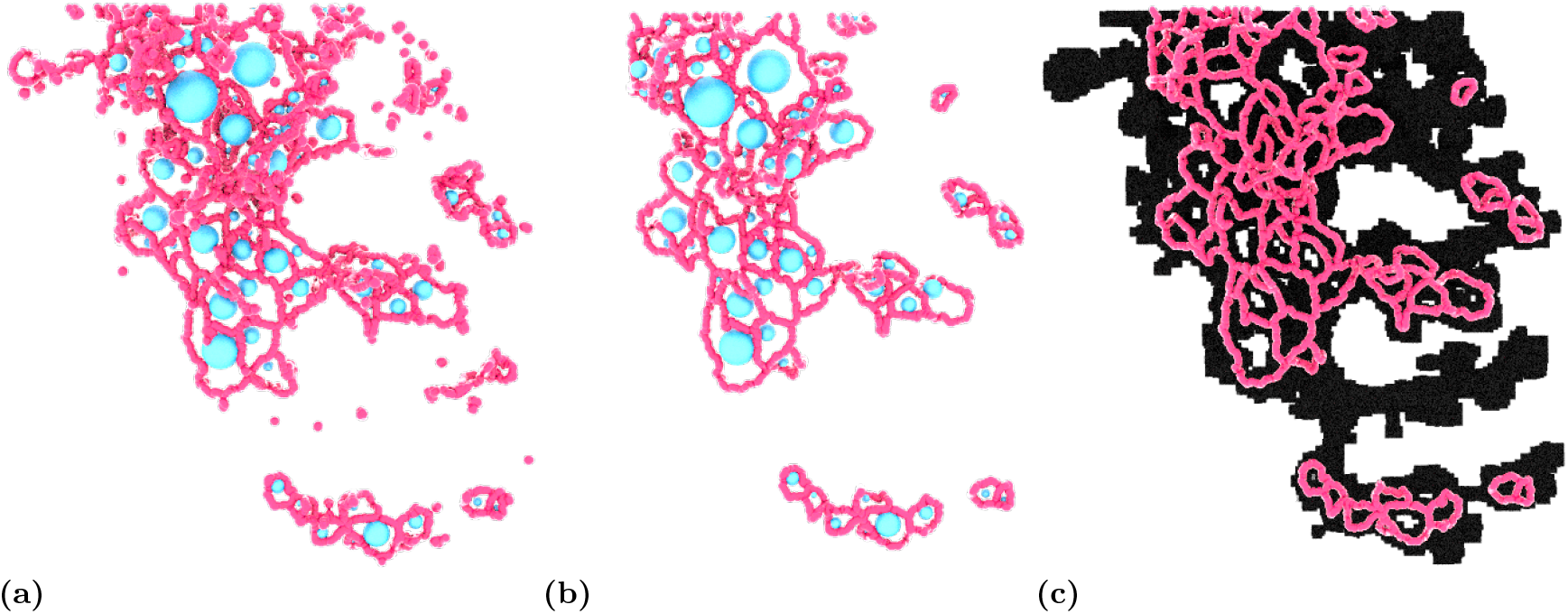
Demonstration of the correspondence between loops and persistence points. Blue spheres indicate the death points of loops in the data. (a) All loops extracted from the skeleton, shown together with persistence points that identify true loops in the data. (b) Only loops paired with points from the persistence diagram, corresponding to true loops extracted from the data. (c) The loops displayed in (b) overlaid with 2D rendering of the raw voxelised data. (note that loops in the z plane will not be visible in this rendering

Such a pairing constructs a set of loops that are matched with points from the persistence diagram and can be used to analyse loop behaviour in the pancreas. By extracting loops via persistence diagrams with geometric representatives, we significantly reduce the number of loops under consideration. For example, depending on the network type, in NEU we retain 47 loops out of 290 present in the skeleton; in DUC, 461 loops out of 5612; and in VAS, 993 loops out of 12 108.

The final step of our analysis is to quantify the “threading” among loops in all the networks. Our approach is inspired by chromatic persistence [12] and is used to study interactions between loops. The main idea is to compare the persistence diagrams of two individual structures with the persistence diagram of their union. Since each point in a persistence diagram represents a loop, changes in the diagram of the union reflect how loops interact when the structures are combined.

To classify loops as threaded or non-threaded, we compare the one-dimensional persistence diagram of a single network with the persistence diagram of its union with another network. Each point in a persistence diagram is represented by a (birth, death) coordinate pair, together with the spatial coordinates of its birth and death voxels.

- **Identification of threaded points**. We identify persistence points from the single-network diagram that do not appear in the persistence diagram of the union. Specifically, we compute the set difference, selecting points whose (birth, death) values are present in the single-network diagram but absent from the union diagram. These points correspond to cycles that become homologically trivial when the second network is added. We interpret such cycles as belonging to the kernel of the induced map on homology and classify them as threaded.
- **Handling multiplicity changes**. When a (birth, death) pair appears in both diagrams but with different multiplicities, we perform an additional spatial consistency check. For each candidate point in the single-network diagram, we verify whether any birth voxel from the union diagram lies within a threshold distance of 5 voxels from the corresponding loop geometry. If no spatially consistent birth is found in the union—indicating that the loop’s birth generator has effectively disappeared—we classify the loop as threaded. Otherwise, the loop is considered unchanged and classified as non-threaded.
- **Classification of geometric representatives**. The homological threading step produces sets of birth and death coordinates corresponding to threaded persistence points. To classify the geometric loop representatives obtained from the pairing procedure, we check whether each loop’s death coordinate (or birth coordinate) lies within a small spatial tolerance (± 1 voxel in each direction) of any coordinate in the threaded set. This fuzzy matching accounts for coordinate rounding differences between the persistence computation and the loop pairing. A loop is classified as threaded if its death point matches the threaded set; otherwise, it is classified as non-threaded.

Figure 5 illustrates examples of threaded and non-threaded loops together with their corresponding persistence diagrams. The mathematical framework of kernel and image persistence was introduced in the form of 6-pack theorem [11], and recently extended to alpha complexes under the name “chromatic persistence” [12]. Although we work with cubical complexes from CT scans rather than alpha complexes, the underlying algebraic foundation, comparing the persistence of a subcomplex to that of the full complex via inclusion-induced maps, is identical. We adopt this terminology and refer to our approach as chromatic persistence throughout the manuscript.

**Fig 5.**
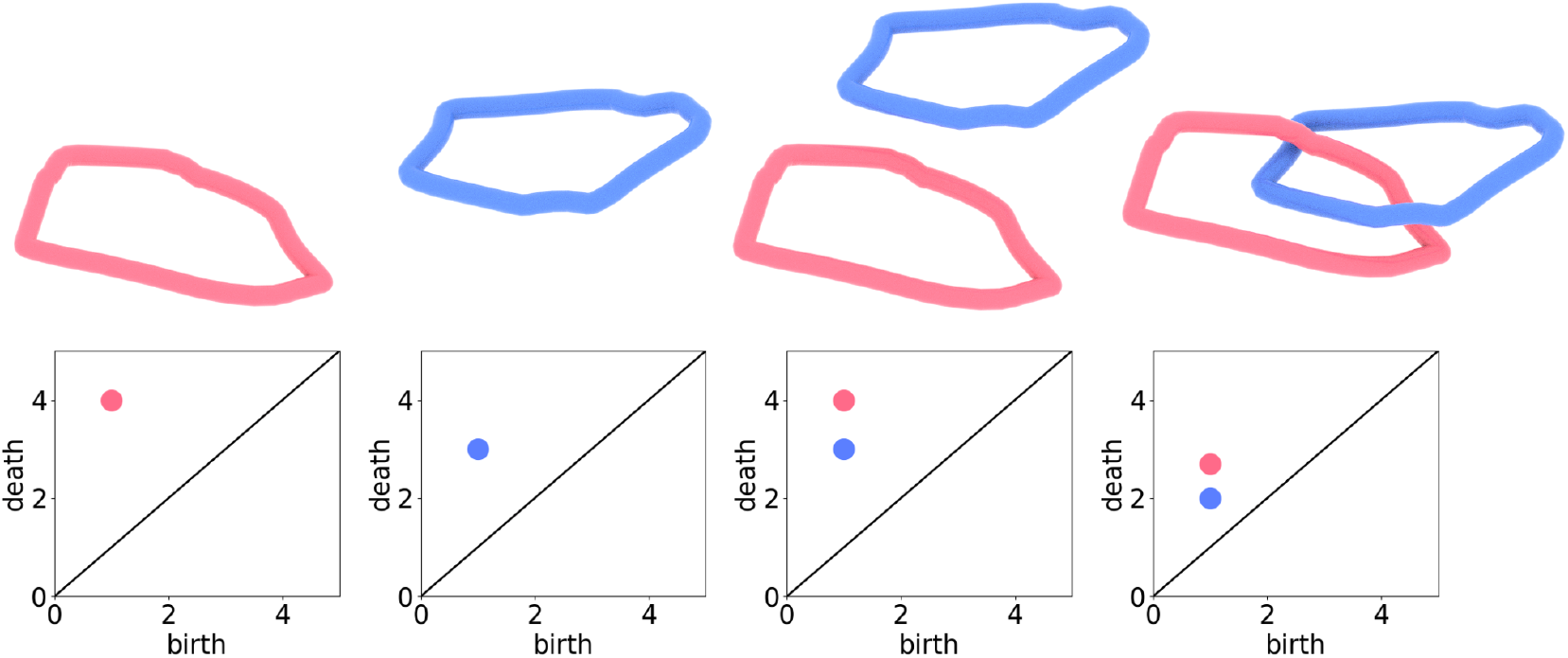
Demonstration of classifying loops through chromatic persistence. Each loop corresponds to a point in the persistence diagram below it. The first two graphics show separate loops, each appearing as a single point in persistence diagrams under SEDT filtration (see Figure 3). The third graphic has two loops that are not linked, and their persistence diagrams remain unchanged, i.e., birth and death do not change for either loop. The final case consists of two linked loops, and their interaction is visible in the change of the death value for both loops: once linked, both cycles are filled at the same filtration value.

**Remark.** In applications to loops, the chromatic approach cannot differentiate between loops that form a topological link and loops that are simply located near each other, as it detects changes in filtration values rather than the exact geometric configuration. However, in both cases, the proximity of loops from different networks reflects a greater degree of entanglement, which is exactly what is relevant for our analysis

## Results

The developing dorsal pancreas undergoes substantial growth between E12.5 and E14.5, during which the ductal, neuronal, and vascular networks expand at different rates and establish increasingly complex spatial relationships (Figure 6). At E12.5, the organ is dominated by the ductal network, while the vascular network is sparse. By E14.5, the vascular network has expanded into a dense network that envelops the ducts. The neuronal network, largely absent at E12.5, where only unconnected cells are found, has developed large-scale loop structures by E14.5. We analyse these changes in two parts: first, we characterise loop formation within each network individually, and then we quantify inter-network entanglement through chromatic persistence.

**Fig 6.**
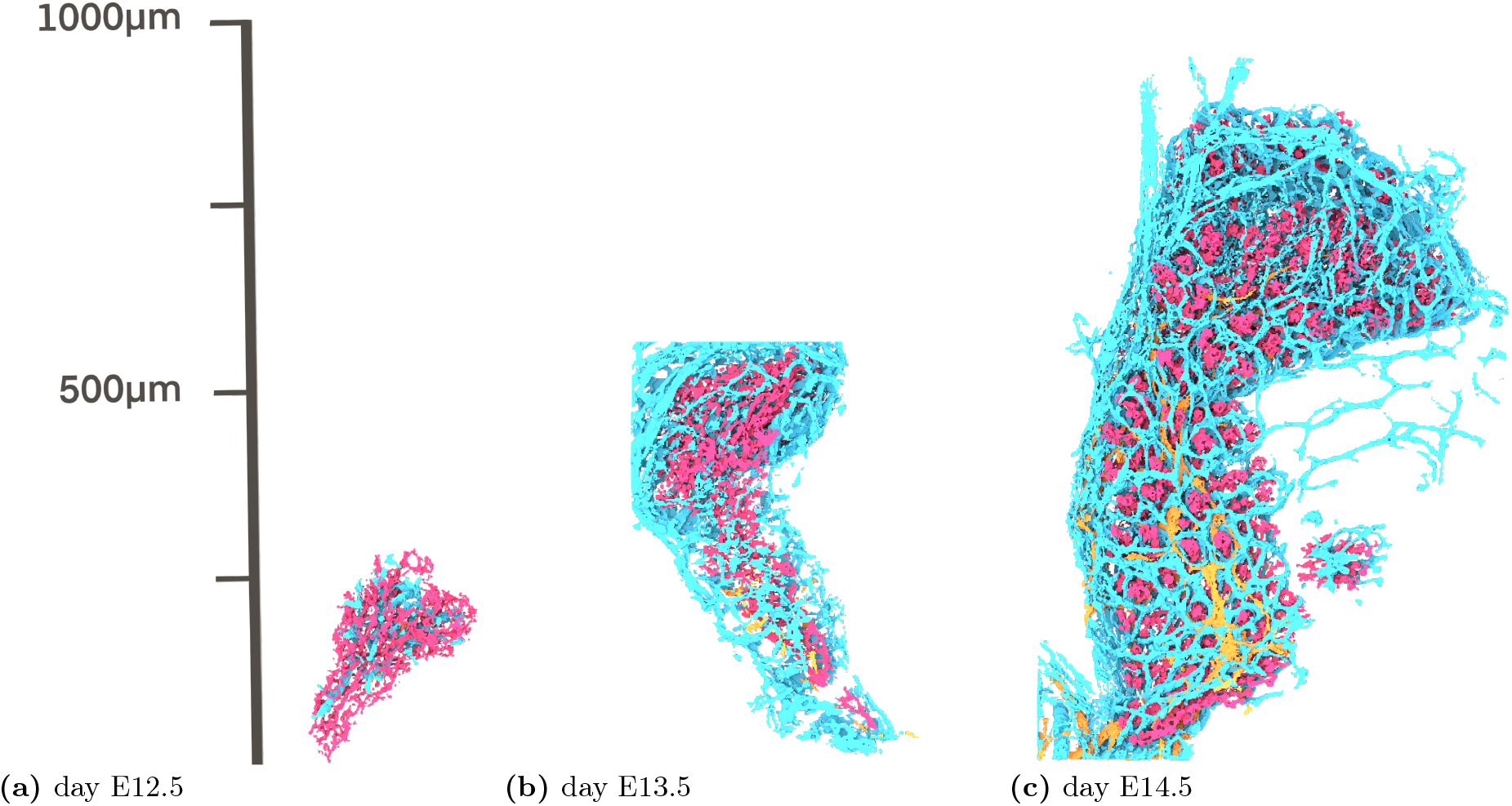
Development of the embryonic mouse pancreas at three stages: (a) E12.5, (b) E13.5, and (c) E14.5. Three tissue networks are shown: neurons (NEU, yellow), ductal epithelium (DUC, pink), and vasculature (VAS, blue). The visualised data is reconstructed from voxelised imaging data, with all three structures present at each developmental stage. At E12.5, the pancreas is compact and dominated by the ductal epithelium. By E13.5, the organ elongates and the vasculature begins to form a more extensive network surrounding the ductal tree. At E14.5, the pancreas has grown with a dense vascular network enveloping the branching ducts and the neural network interspersed throughout. Images are shown at their original scale in *µm*, highlighting the growth of the organ across developmental stages.

We provide general statistics and insights into network threading during dorsal pancreas development. Specifically, we present the total number of loops and how this quantity changes throughout development. We then analyse changes in loop size distributions over developmental time. Next, we examine the spatial distribution of loops across the entire organ and explore the relationship between loop size and spatial location.

In the second part of this section, we focus in more detail on threading behaviour, primarily investigating interactions of DUC with NEU and VAS, as the entanglement of the ductal network with each of the other two networks is of particular interest for understanding pancreas development. We quantify how threading evolves throughout pancreas development and analyse which regions of the organ exhibit more active threading. We consider developmental stages at E12.5, E13.5, and E14.5 days, with 5, 3, and 3 samples from different embryos, respectively. Figure 6 demonstrates the raw data of all three networks all together at different stages of development. Figures 7, 8 and 9 visualise each network and its development.

**Fig 7.**
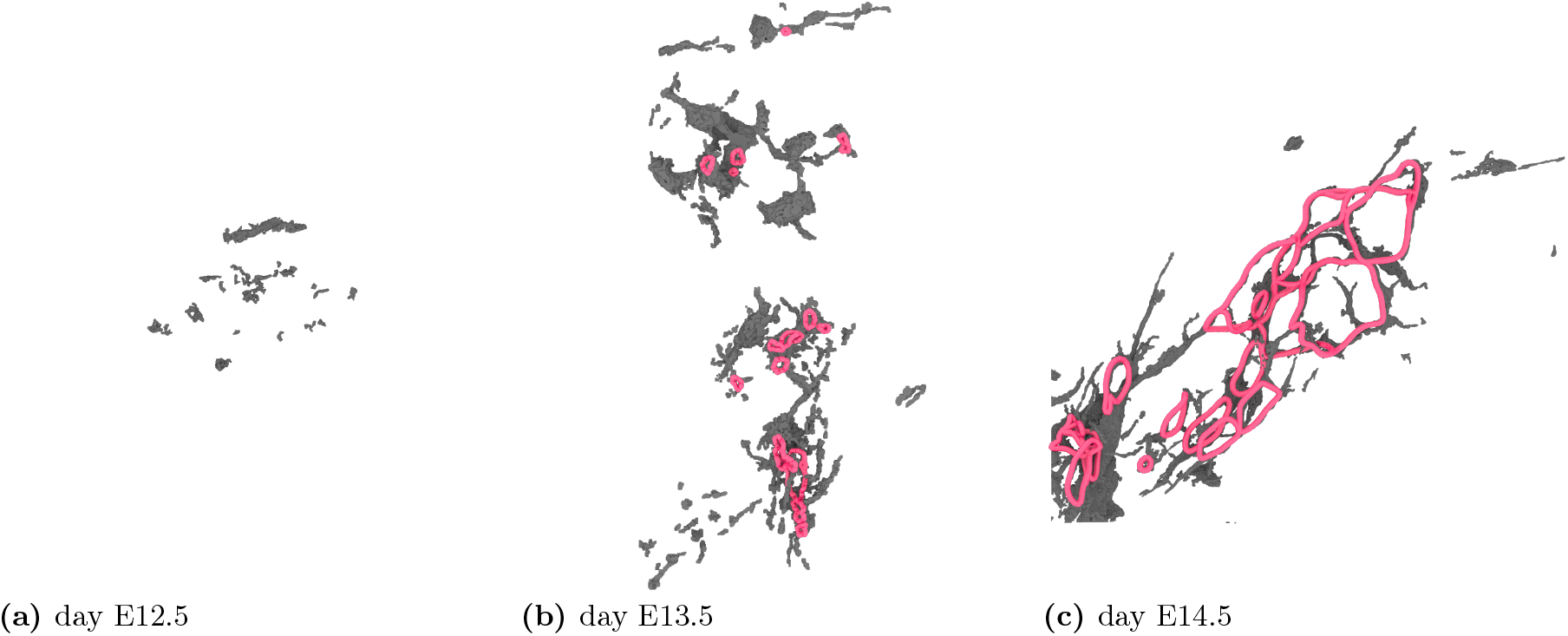
Demonstration of the number of loops present in the neural (NEU) network at each developmental day. The pink colored structures correspond to extracted loops, and the black outline is raw voxelised data. The scale is preserved to demonstrate the scale of development.

**Fig 8.**
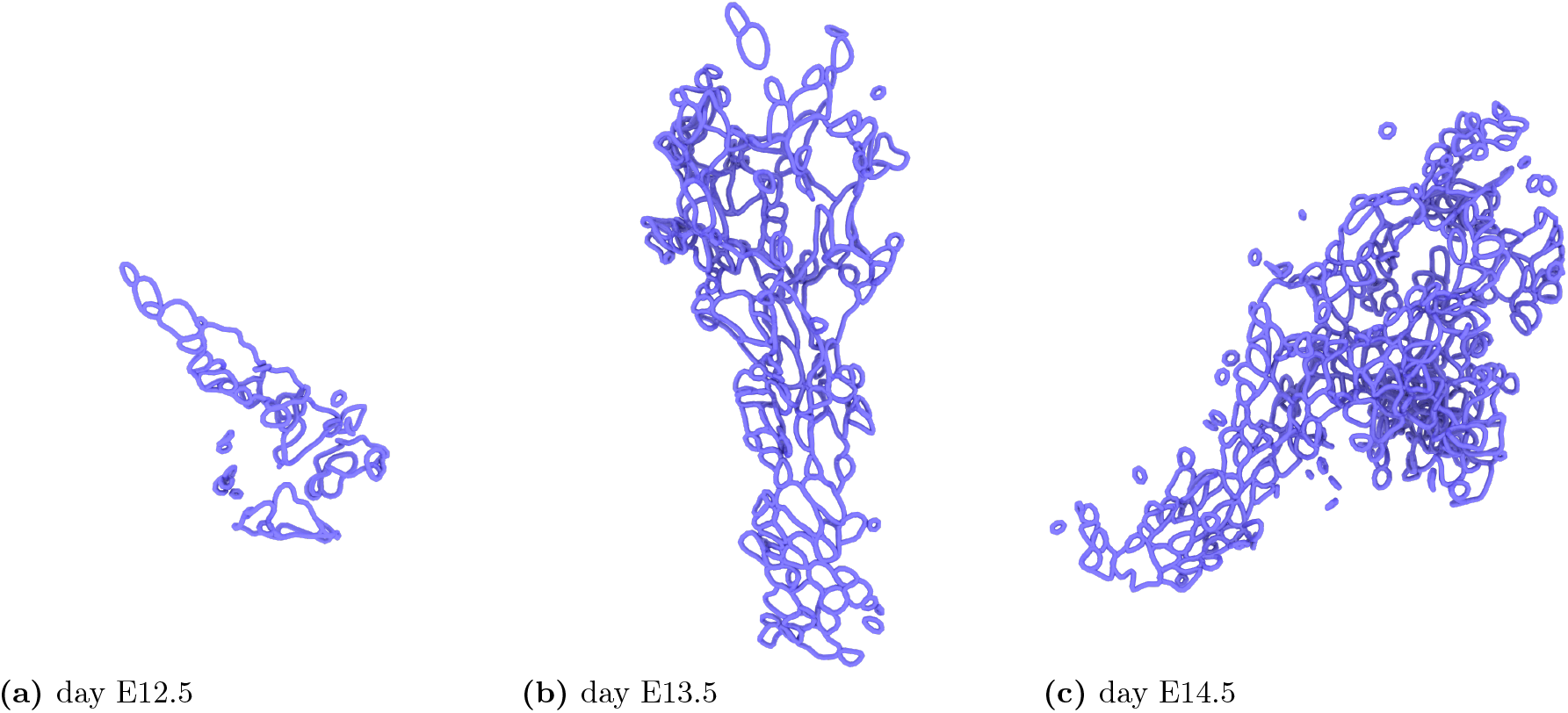
Demonstration of the number of loops present in the ductal (DUC) network at each developmental day. The scale is preserved to demonstrate the scale of development.

**Fig 9.**
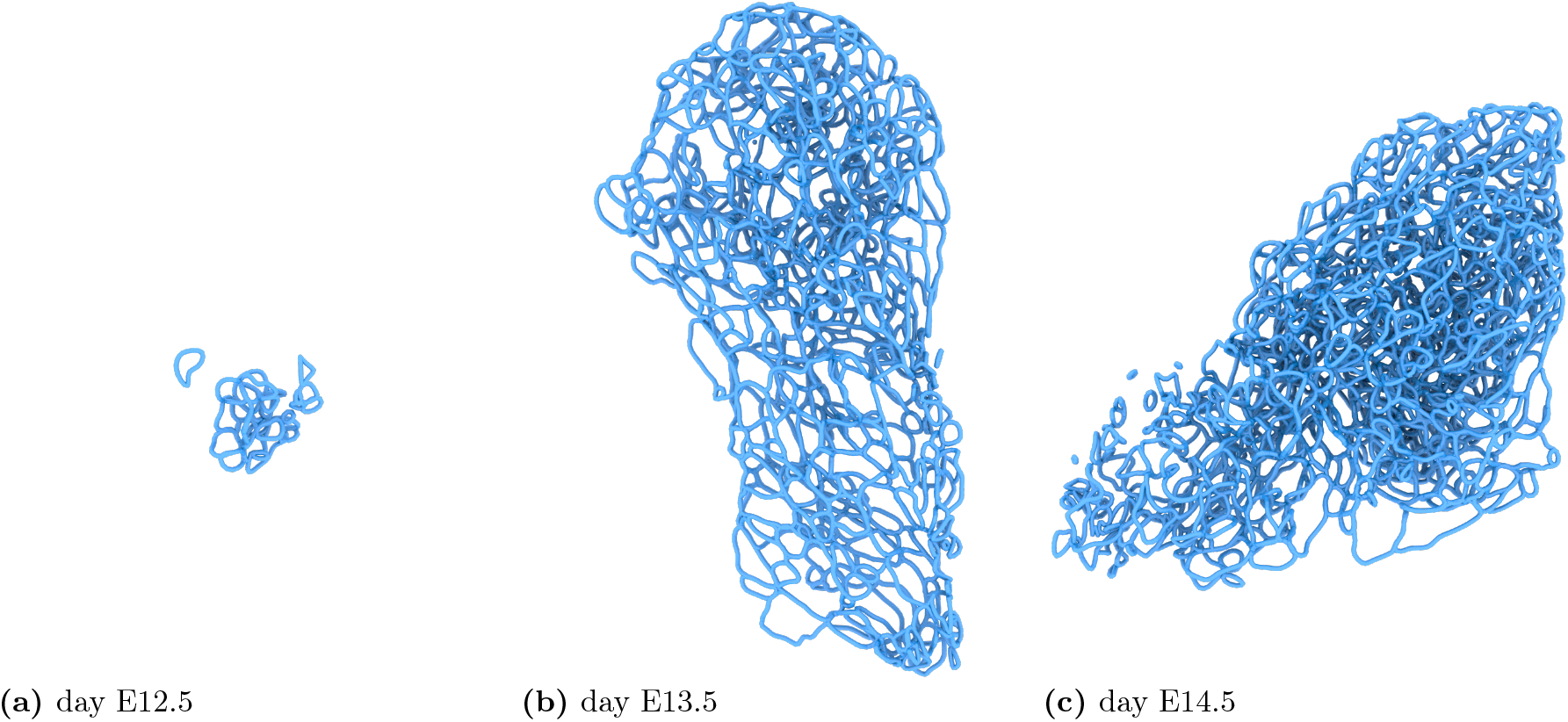
Demonstration of the number of loops present in the vascular (VAS) network at each developmental day. The scale is preserved to demonstrate the scale of development.

### Temporal and spatial separate analysis of neural, ductal, and vascular networks in the pancreas

#### Loop development

Loop formation does not start at the same time across all networks. In the neural network(NEU), loops first appear at E13.5. By E14.5, the network contains large-scale loops, see Figure 7. In the ductal network(DUC), loops are already present from day E12.5. Figure 8 shows an example of loop development in DUC. Finally, loops in the vascular network (VAS) start forming as early as day 12.5 and surpass those in DUC in total count at later stages (Figure 9). Figure 10a shows the total number of loops over developmental time for all networks. We observe that the number of loops in DUC and VAS on day E13.5 lies within a smaller range than on day E14.5; in particular, the loop counts in DUC and VAS on day E14.5 differ substantially.

**Fig 10.**
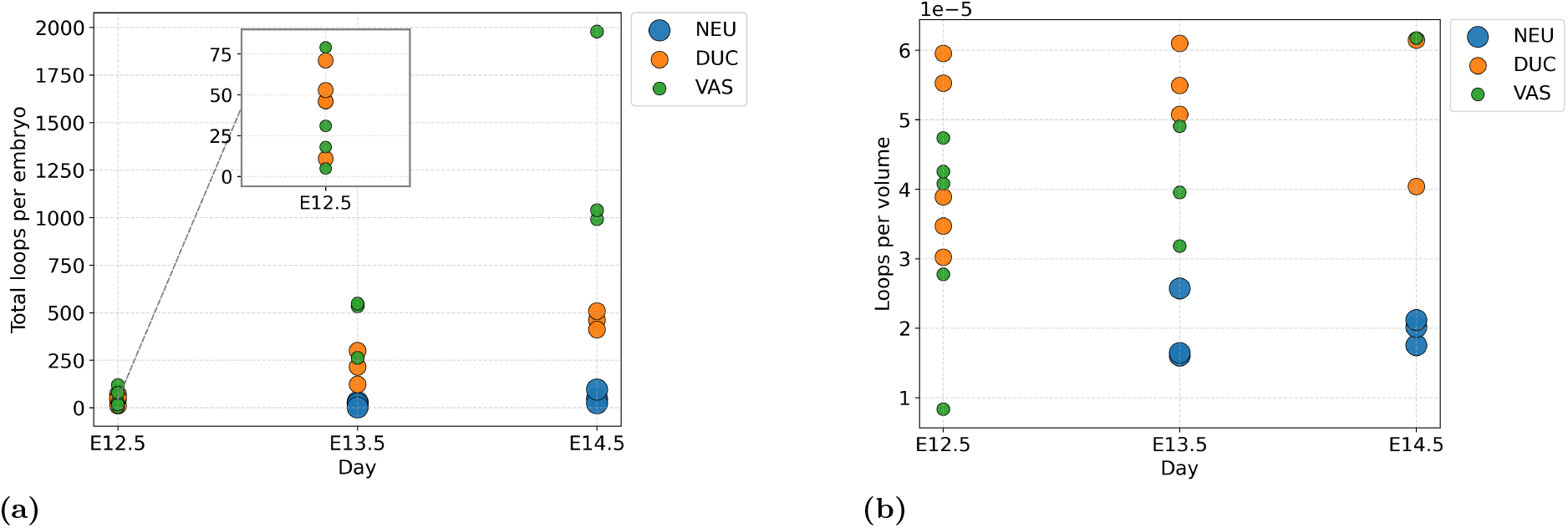
Total counts of loops. (a) Number of loops in each type of network for each embryo. (b) Number of loops in each type of network for each embryo, normalised by the volume of voxels in the network.

Another way to quantify loop development during growth is to consider the number of loops normalised by volume of the segmented network (Figure 10b). The relative loop density in VAS shows a slight upward trend over development, indicating a tendency away from a tree-like structure. In contrast, DUC shows a decrease in relative loop density from day E13.5 to day E14.5.

#### Loop sizes

Figure 12 shows the distribution of loop sizes in each network type across developmental stages. By the size of a loop, we mean curve length. The peak of the distribution for NEU is at 70.4 *µ*m at day E14.5 and 43.7 *µ*m at day E13.5. However, the distribution of the loop size demonstrates that there are only a few large loops and most of the detected loops are less that the peak value of 43.7 *µ*m. For DUC, the peak across all stages is at 66.5 *µ*m. For VAS, the peak is at 63.2 *µ*m at day E12.5 and shifts to 81.1 *µ*m at days E13.5 and E14.5. Figure 11 shows examples of loops in each network, coloured by size.

**Fig 11.**
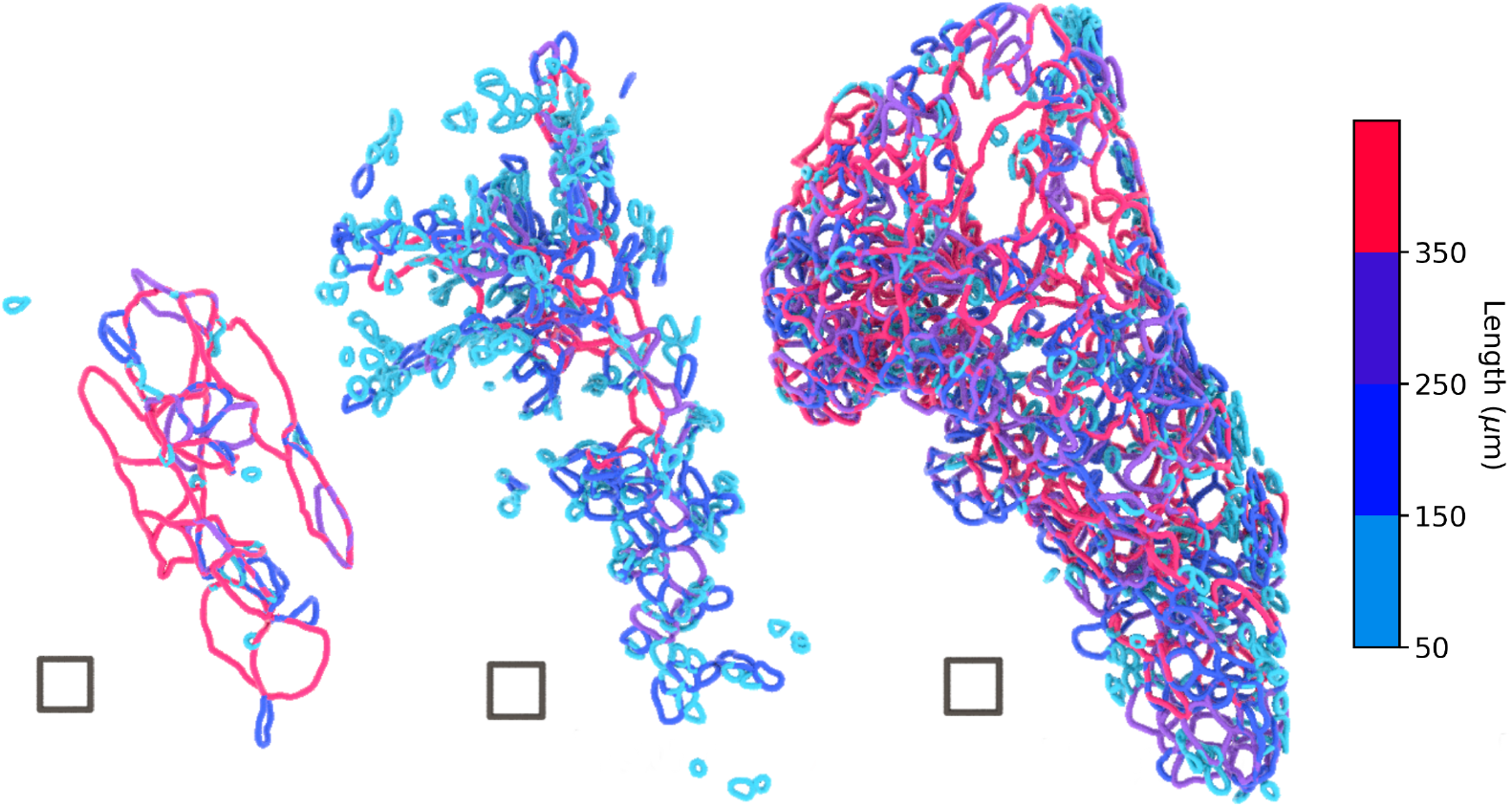
All three networks Neurons(left), Ducts(middle), and Vasculature(right) are from the same embryo at day E14.5 and respective loops are colored by size. The square has each side 50*µm*.

**Fig 12.**
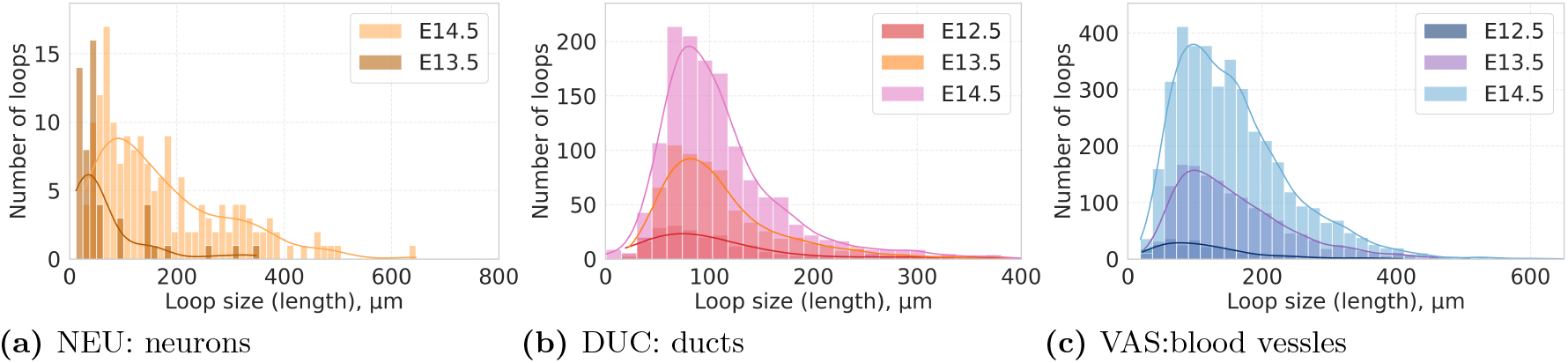
Loop size distributions for each network type across all embryos. Each plot shows separate histograms for different developmental days. (a)Neural network. No loops are present at day E12.5; therefore, only days E13.5 and E14.5 is shown. (b) Ductal network. (c) Vascular network. Note that the number of loops varies substantially between network types, and thus the axes are not normalised.

We observe that the structure of DUC consists of smaller loops than NEU, even though the peaks of the histograms differ only slightly (approximately 14 *µ*m). We also observe that the size distribution at day E13.5 already appears very similar in shape to that at day E14.5.

#### Loop distribution

The death points from persistence diagrams provide the spatial locations of loops, allowing us to study how loops in DUC and VAS are distributed throughout the pancreas. Pairing persistence points with geometric representatives tends to undercount loops; since we do not require geometric representatives here, we instead use the coordinates of death points taken directly from the persistence diagrams, after cleaning to retain unique death points.

We estimate the density distribution and visualise 2D slices taken from the middle of the volume, as well as 2D slices taken 20 planes before and after the centre. The intensity scale is adjusted separately for each figure to better display the density level sets; this sacrifices a uniform scale across all networks and developmental stages.

We also plot a schematic 3D histogram, in which we count the number of loops (death points) within cubic bins of side length 60 *µ*m. Spheres are plotted at the bin centres, with their radii proportional to the number of loops within each bin. Figure 13b and 13d show one sample on E14.5, and one on day E13.5. Due to the small sample size, we cannot draw statistically significant conclusions about uniformity; however, we observe that in some samples, loops are more concentrated in central regions. Similarly, in VAS the loop density is higher towards the centre (Figure13f and 13h). The plots for all the samples can be found in the Supplementary Information.

**Fig 13.**
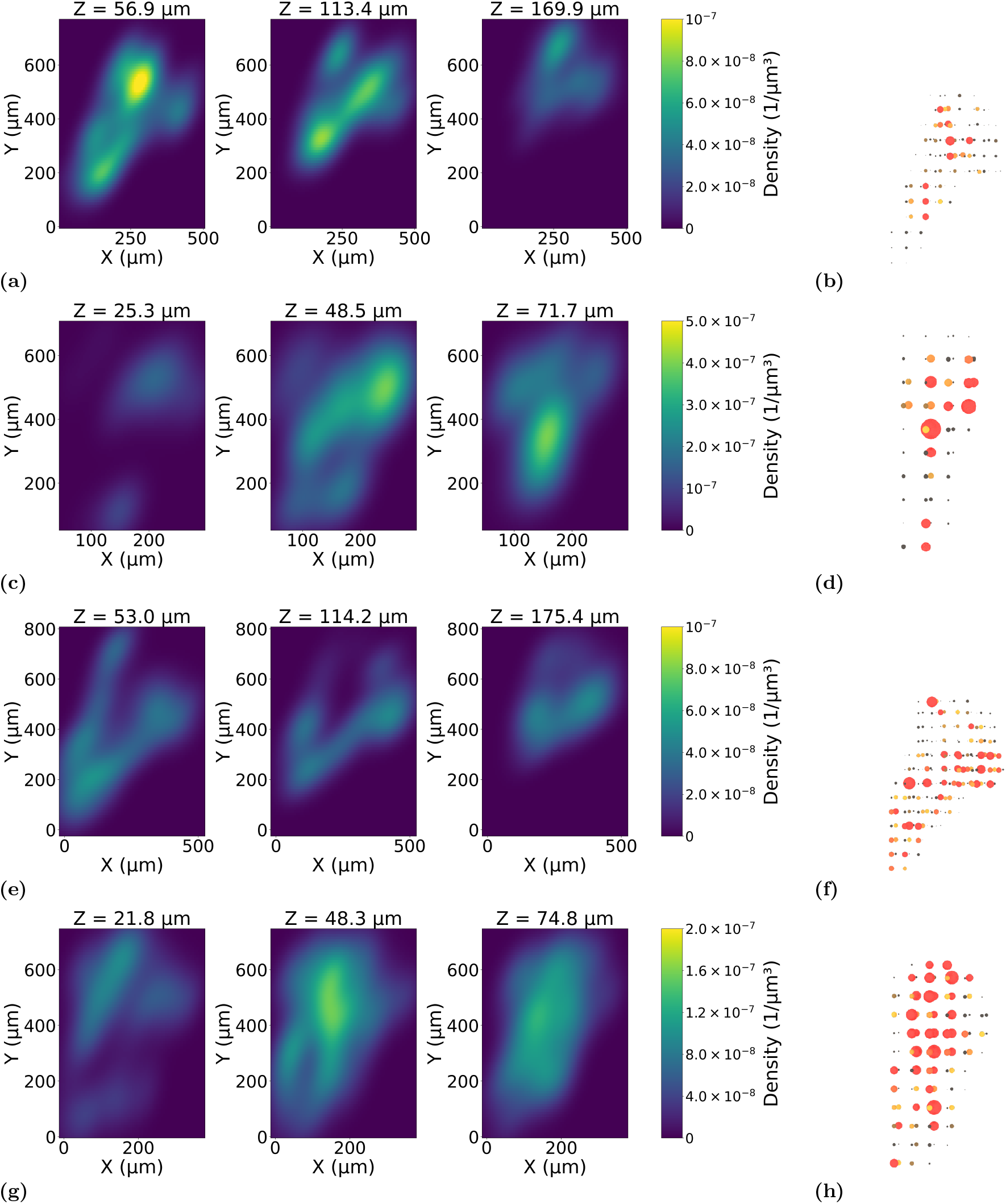
Density of loop distributions in DUC and VAS. In each row, the middle slice is taken at the centre of the volume along the shortest axis (*z*), while the left and right slices are taken at ±20 slices. The intensity scale is not kept constant across samples to better illustrate the distributions. Subfigures (a)–(b) correspond to DUC at day E14.5 of development, (c)–(d) at day E13.5, (e)–(f) to VAS at day E14.5, and (g)–(h) at day E13.5. The subfigures in the leftmost column show a schematic three-dimensional histogram: bins correspond to cubes of side length 60 *µ*m, and each sphere’s radius represents the number of loop centres (death points) in the corresponding cube. Colour highlights regions with high concentration of loops, with red indicating the highest densities.

We also wanted to observe if there is any correlation between the loop size and its locations. We therefore build a mask that envelops the union of networks by inflating the networks and closing the holes. Then we build a distance map slice-wise that evaluates the distance from each voxel to the boundary of the enveloping mask, see Figure 14

**Fig 14.**
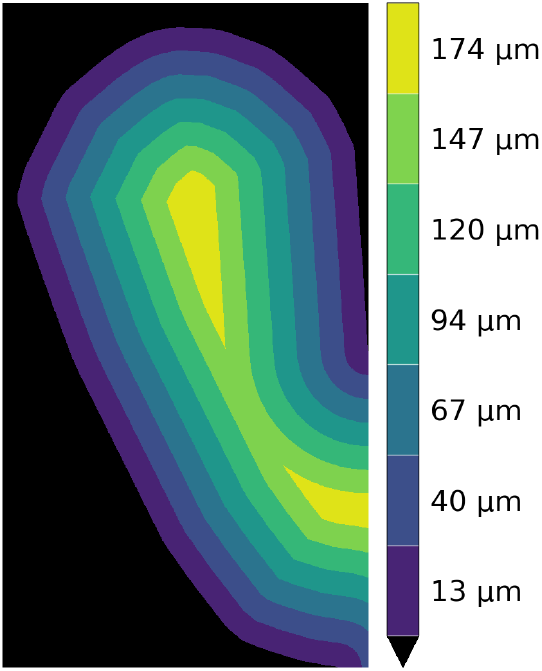
An example Discretised levelsets of the distance map of the enveloping mask. The numbers on the bar show the value of the distance in the middle of each level set.

**Fig 15.**
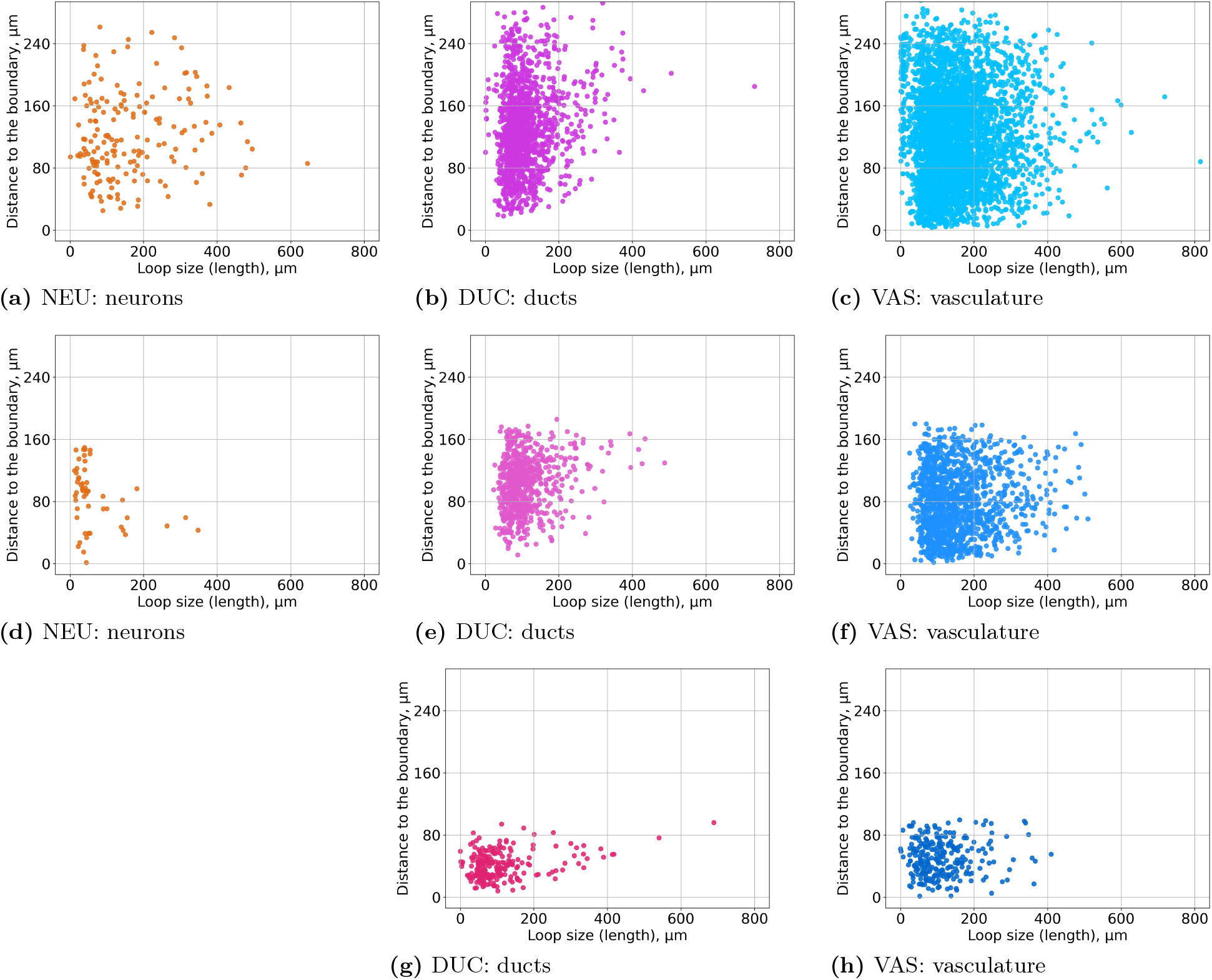
Loop size vs distance of the death point to the enveloping area of all the networks. Each row corresponds to a different day of development: (a)–(c) E14.5, (d)–(f) E13.5, (g)–(h) E12.5.

For each loop, we look at the distance from the border to the paired death point. We opt for the death points because they are nearly guaranteed by the pairing algorithm to be spatially well-localised with respect to the loop. In contrast, the geometric barycentre of a highly non-convex loop can lie far from the loop itself. And we plot the distance from the death points to the border against the size of loops. For neurons and vasculature we observe no correlation between size and position. For ducts, we also observe no linear correlation, but do observe that large loops are always furthest from the border (i.e. in the centre of the pancreas), which confirms an impression given by the visualisation, Figure 11.

#### Threading analysis of neural, ductal, and vascular networks in the pancreas

In this section, we quantify network interactions through their threading and entanglement. All three networks grow together in the pancreas while being intertwined (see Figure 16). They grow through each other at different rates: as shown in Figure 10a, on day E12.5, the ductal network DUC has, on average, more loops already formed than the vasculature VAS, but VAS surpasses DUC on day E13.5, and the difference in loop number is quite large on day E14.5. We focus on exploring the entanglement of the ductal network with vasculature and neurons through loop interactions using geometry. Importantly, the loop structure in the networks is quite dense (Figure 16); therefore, describing loop interactions captures much of the information about the relationships between the networks as full structures.

**Fig 16.**
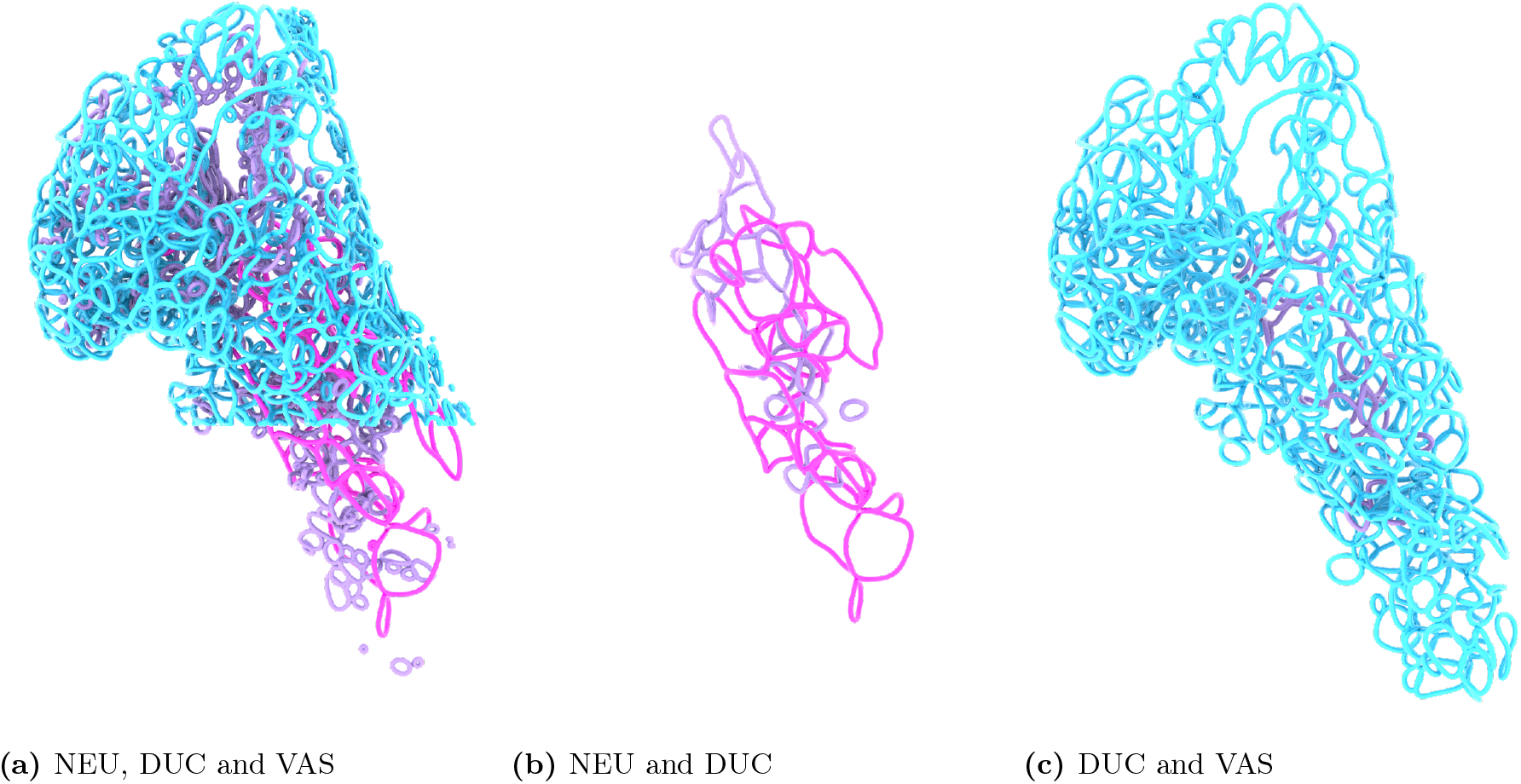
(a) All loops extracted from the three networks shown together: NEU (pink), DUC (purple), and VAS (blue). Only loops are displayed, not the full network skeletons. The vascular network is partially cut away at the bottom to expose the spatial relationship between DUC and NEU. (b) DUC entangled with NEU, i.e., only loops that are classified as threading for NEU via DUC and for DUC via NEU. (c) DUC entangled with VAS, i.e., only loops that are classified as threading for VAS via DUC and for DUC via VAS.

Using the persistent homology calculations, we quantify the number of threaded and non-threaded loops. The networks consist of loops and branches; therefore, in our approach, we have three geometric configurations when chromatic persistent homology detects threading: Two loops form a topological link, two loops intersect or approach each other without forming a link,and one loop is threaded by a non-loop segment, see Figure 17. Threading can always occur via two other networks that are distinct from the considered; therefore, we report the cases based on which network the thread occurs. We do not report all the possible combinations but rather those that are of particular significance.

**Fig 17.**
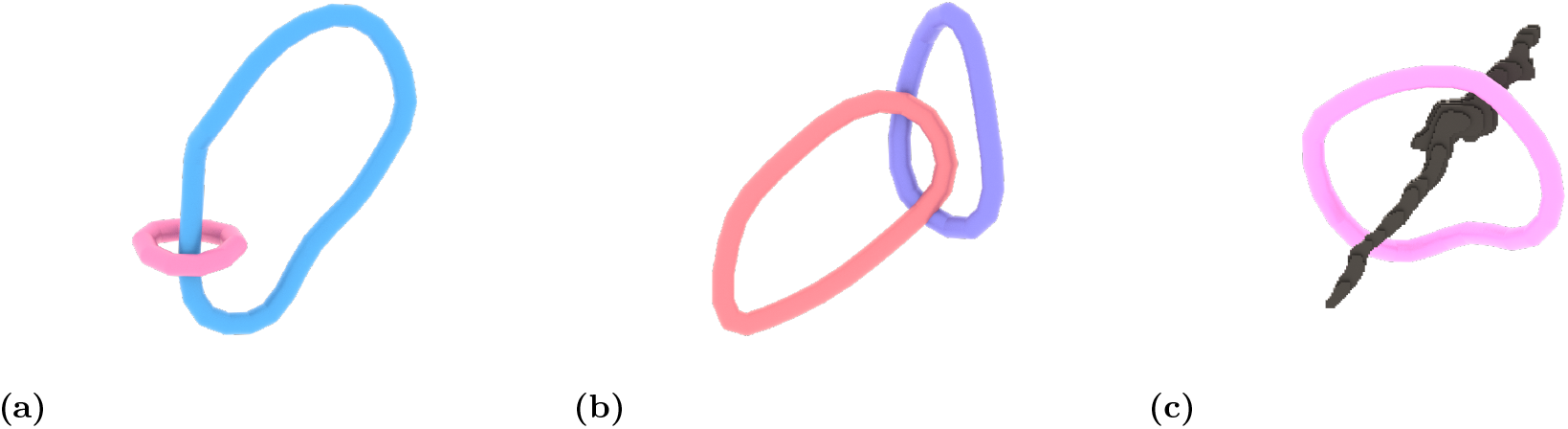
Demonstration of threading configurations. We demonstrate possible cases on the loops taken from the data E14.5. (a) Two loops form a topological link. (b) Loops intersect or approach each other without forming a link. (c) A loop is threaded by a network segment that does not belong to any loop.

The neurons (NEU) only have fully formed loops at E14.5 days, and they can be threaded via DUC or VAS, as shown in Table 1. We observe that the neural network NEU is entangled with both networks, but more entangled with the vascular network VAS, with on average 84% being threaded; however, about 58% of loops are entangled by the ductal network DUC on average.

**Table 1.** The number of loops threaded in NEU (neurons) via DUC (ducts) and VAS (vasculature) per different embryo on day E14.5. In brackets, we report the percentage of threaded loop in each sample.

|  | Emb1 | Emb2 | Emb3 |
| --- | --- | --- | --- |
| DUC | 29 (62%) | 15 (60%) | 51 (53%) |
| VAS | 33 (70%) | 23 (92%) | 86 (90%) |

If we compare the shape of the distributions in Figure 18 to the distribution in Figure 12a we see that there are significantly fewer small loops, which leads us to the conclusion that larger loops in NEU are entangled by DUC, but smaller loops are left unthreaded by DUC. However, we can see that almost all loops are threaded with VAS and only a small number of smaller loops are left unthreaded.

**Fig 18.**
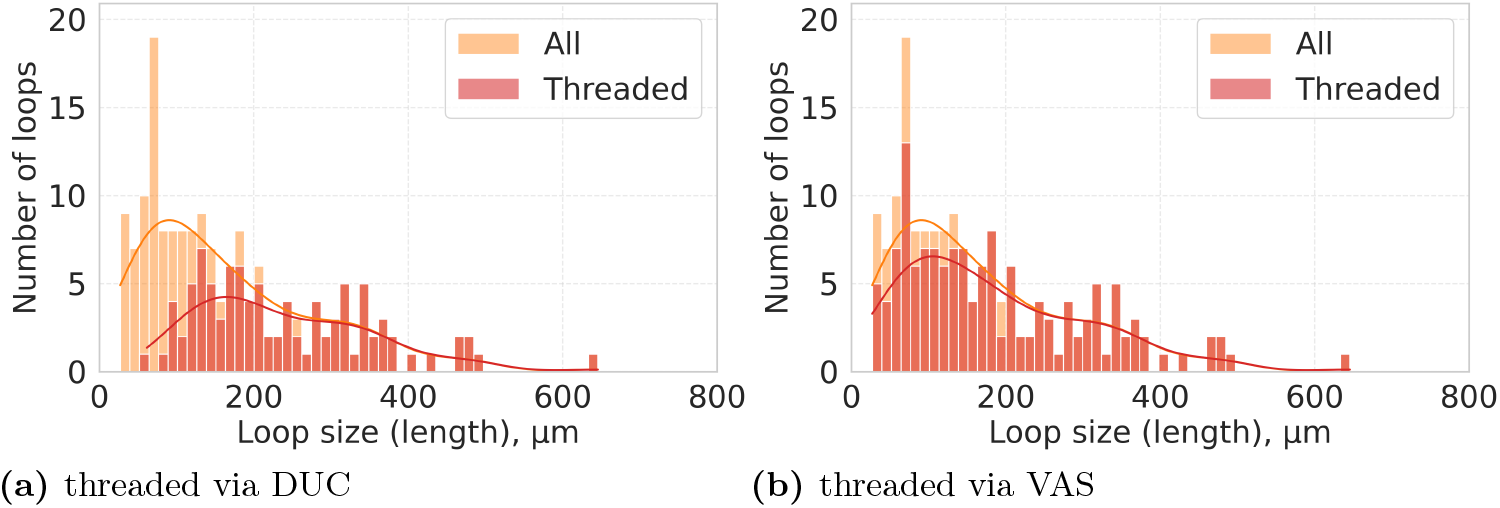
Size distribution of threaded loops in NEU. The orange colour demonstrates the histogram of all loop sizes(length) over all samples. The pink histogram corresponds only to threaded loops.

In both DUC and VAS, loops are present at all developmental stages, allowing us to track threading over time; the total number of threaded loops for each case is presented in Figures 19a and 19b.

**Fig 19.**
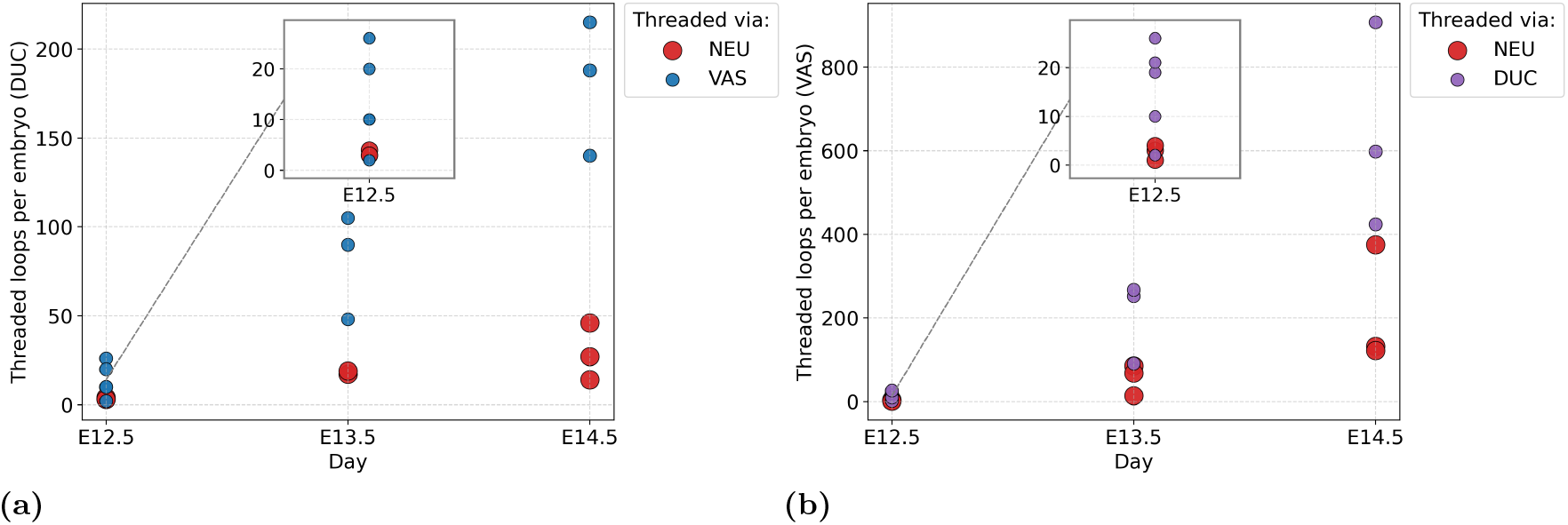
Total counts of threaded loops, where the legend indicates which network is threading. (a) The total number of threaded loops in DUC via NEU and VAS for each day and embryo. (b) The total number of threaded loops in VAS via NEU and DUC for each day and embryo.

First, we look at the percentage of threaded loops in DUC via NEU and VAS (see Figure 21a). There is not much variance between different embryos, especially on days E13.5 and E14.5. As we can see, although the absolute number of threaded loops via NEU slightly increases (Figure 19a), the proportion of threaded loops remains almost the same, 5%± 4%, when averaged over all days and embryos. The threading of DUC via VAS is much higher and changes over the days: 28% ±12% (E12.5), 36% ±2% (E13.5), and 40%± 11% (E14.5). We observe that the later days have a small variance among different embryos.

In the case of VAS, the variance in threading values on day E12.5 is quite large (≈20–30%) (Figure 21b); however, for the later days, we can present meaningful averages. The threading of VAS via NEU stays quite low and increases as the network grows: 3% ±4% (E12.5), 11% ±5% (E13.5) and 15%± 4% (E14.5). Interestingly, the threading of VAS via DUC increases, as in the DUC via VAS case, with averages of 42% ±18% (E12.5), 39%± 6% (E13.5) and 50% ±7% (E14.5).

If we look at the number of threaded loops in DUC and VAS normalised by the volume of the whole structure (Figure 20), we see that the behavioural trend remains similar to the one presented for the total number of loops (Figure 20). This suggests that the threaded-loop subset evolves in line with the overall loop population, rather than showing a clearly different scaling with network growth over the observed time period.

**Fig 20.**
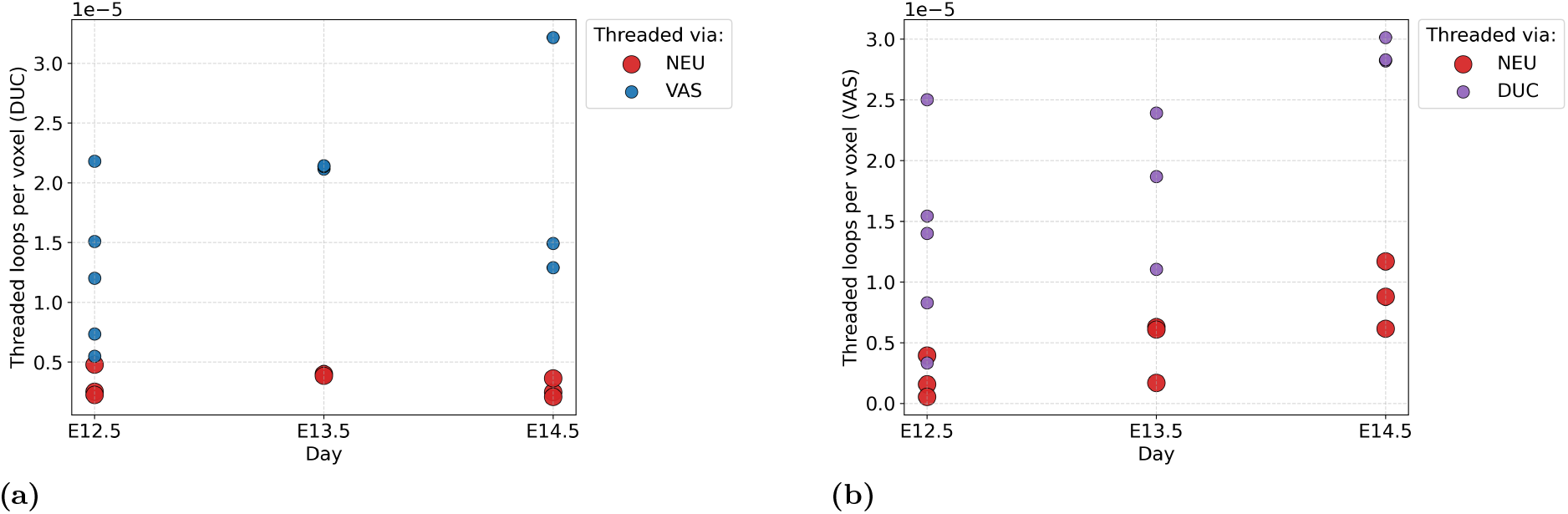
(a) Total number of threaded loops in DUC via NEU and VAS normalised by the volume of the network.(b) Total number of threaded loops in VAS via NEU and DUC the volume of the network.

**Fig 21.**
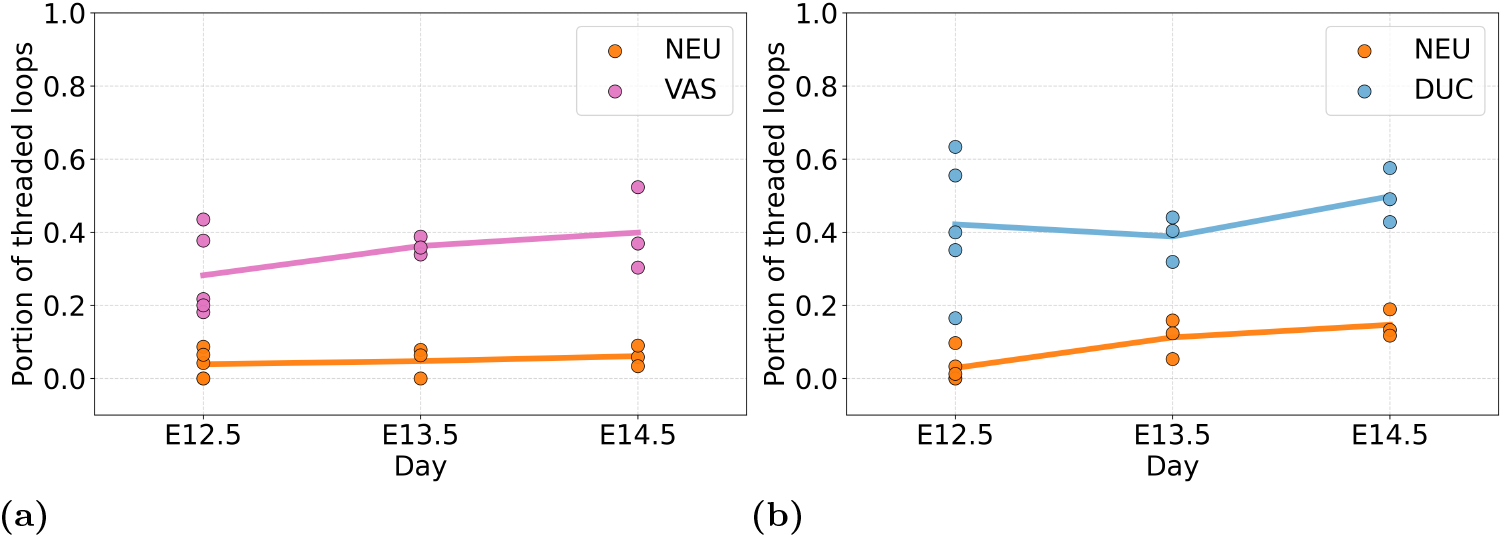
The percentage of loops threaded in (a) DUC via NEU(orange) and VAS (pink) and (b) VAS via NEU(orange) and DUC (blue) throughout the days E12.5, E13.5, and E14.5. The line plot goes through the average point for each day.

We examine threading behaviour in greater detail in the ductal network DUC, as previous work supports the idea that ductal loops play a particularly important role in endocrinogenesis [2]. The pancreatic ductal system is a three-dimensional tubular network whose topology (branching and loops), geometric properties (diameter, curvature, and length), and spatial embedding constrain cellular differentiation and tissue interactions. Quantifying this geometry allows developmental processes to be described in terms of measurable rules rather than purely qualitative observations. Here, we extend these ideas by quantitatively analysing the geometry of the entanglement of the ductal network with neurons and vasculature.

The size distribution of loops in DUC that are threaded by VAS is shown in Figure 22. The peak of the distribution shifts towards larger loop sizes over development, with peak values of 110.3 *µ*m at day E12.5, 95.7 *µ*m at day E13.5, and 81.1 *µ*m at day E14.5. Overall, the distribution is skewed towards larger loops, indicating that larger ductal loops are more likely to become threaded by vasculature, which is an interesting result, given that we have previously shown that larger loops are more stable and have a higher density of beta cells [1]. The size distribution of loops in DUC that are threaded by NEU can be found in the Supporting Information.

**Fig 22.**
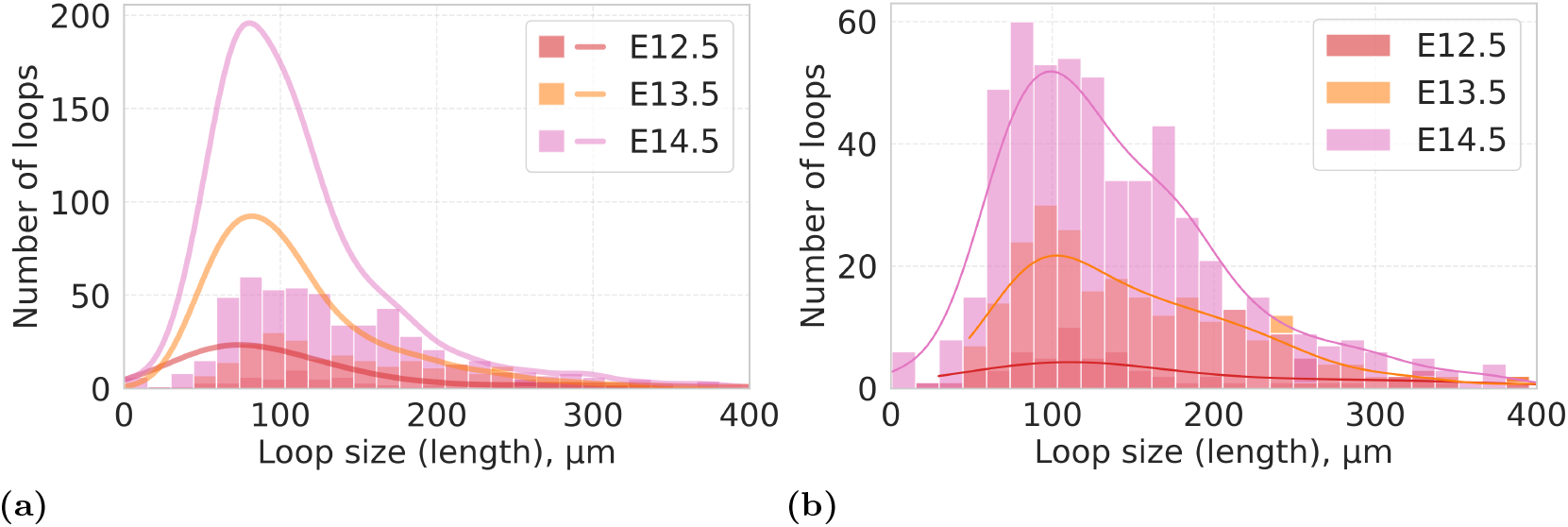
Distribution of threaded loop size in DUC via VAS. (a) The histogram displays the distribution of threaded loops on each day. The approximation of the distribution plotted as lines corresponds to the distributions of the sizes of all loops for each day, respectively. The colors of the lines correspond to the colors of the histograms for each day accordingly. (b) The same histogram as in (a) but on a different y-axis scale for better display of the shapes of the histograms with respect to each other. In both cases, each plot is built over all embryos for each day.

Another aspect of threading is the spatial location of threaded loops. We examine the distribution of loop positions (death points) with respect to their relative depth within the organ, defined as the distance of each loop centre to the organ boundary, normalized by the maximum distance (i.e., the deepest point) within the organ.

We consider the threading of the ductal network, threaded via neurons and the vascular system. In the case of entanglement via the neural network, even though we have a low amount of threading, we can see that it happens more centrally in the organ (Figure 25), which coincides with the visual intuition coming from the data. When it comes to threading via the vascular system on all days, we observe a slight shift in which loops closer to the centre of the organ are more likely to be threaded, while loops located near the boundary tend to remain non-threaded (Figure 25). Since there is no correlation between loop size and spatial position (Figure 15), we conclude that threaded loops tend to be located closer to the centre of the pancreas, which is interesting given our previous finding that beta cells are enriched in the centre of the pancreas [1].

The size distribution of loops in VAS threaded by DUC is shown in Figure 23. The peaks of the distributions occur at 99.0 *µ*m on day E12.5, 170.4 *µ*m on day E13.5, and 152.2 *µ*m on day E14.5. Overall, the distribution is skewed towards larger loop sizes, indicating that larger loops are more likely to be threaded. The size distribution of loops in VAS that are threaded by NEU can be found in the Supporting Information.

**Fig 23.**
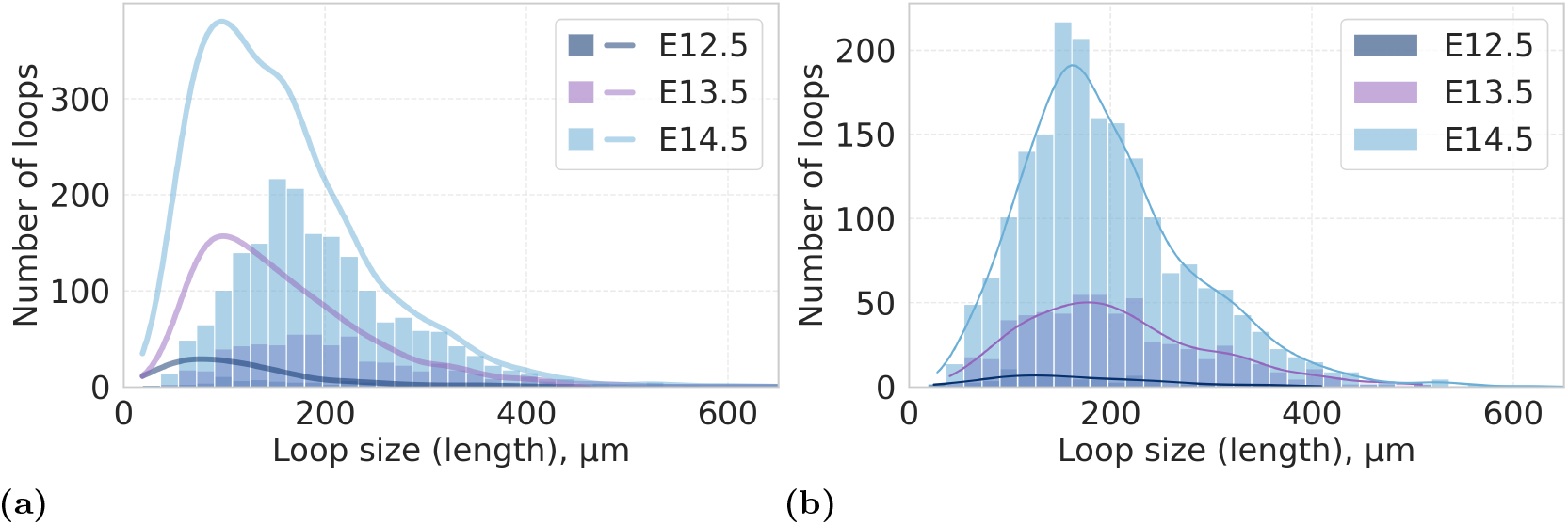
Distribution of threaded loop size in VAS via DUC. (a) The histogram displays the distribution of threaded loops on each day. The approximation of the distribution plotted as lines corresponds to the distributions of the sizes of all loops for each day, respectively. The colors of the lines correspond to the colors of the histograms for each day accordingly. (b) The same histogram as in (a) but on a different y-axis scale for better display of the shapes of the histograms with respect to each other. In both cases, each plot is built over all embryos for each day.

**Fig 24.**
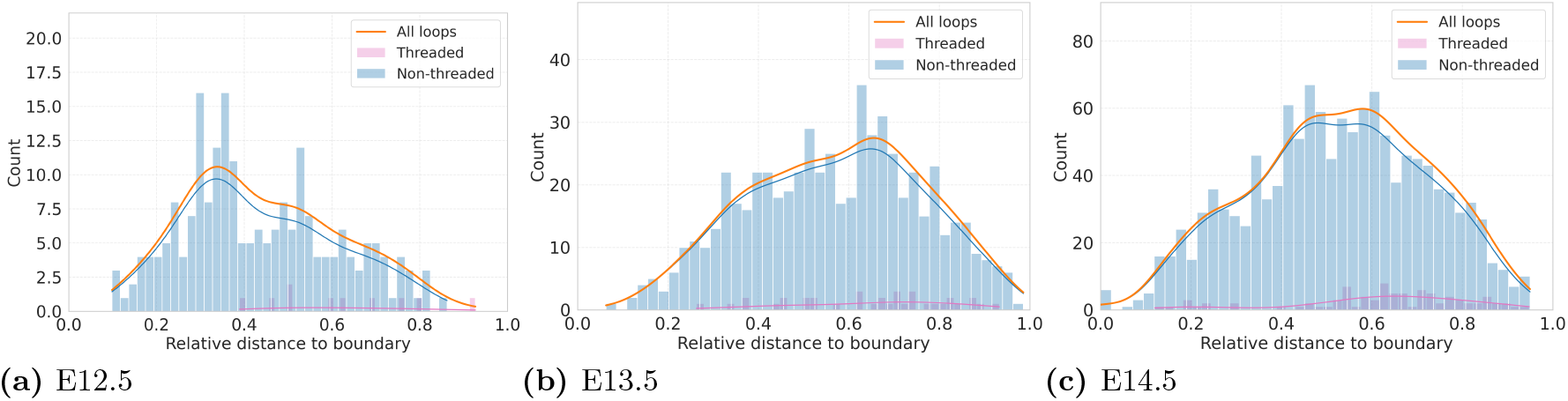
Distribution of the relative position of the death point related to each loop inside the pancreas in DUC. The orange line is the distribution built over all points, while pink corresponds to only the threaded loop via NEU, and blue to non-threaded. The plots are built over all the embryos separately for each day. The histograms do not have the same y-axis to emphasise the shape within each day.

**Fig 25.**
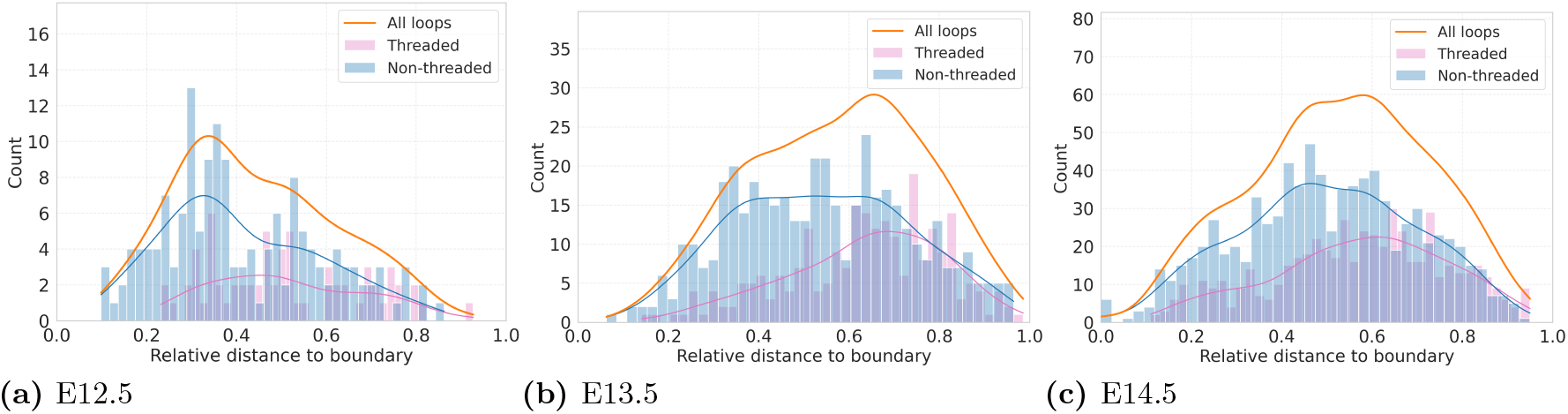
Distribution of the relative position of the death point related to each loop inside the pancreas in DUC. The orange line is the distribution built over all points, while pink corresponds to only the threaded loop via VAS, and blue to non-threaded. The plots are built over all the embryos separately for each day. The histograms do not have the same y-axis to emphasise the shape within each day.

The spatial distribution of threaded loops shows that loops located near the periphery of the organ tend to remain non-threaded, while loops closer to the interior are more frequently threaded (Figure 26). As there is no correlation between loop size and spatial position, we conclude that threading of VAS by DUC predominantly occurs near the centre of the pancreas.

**Fig 26.**
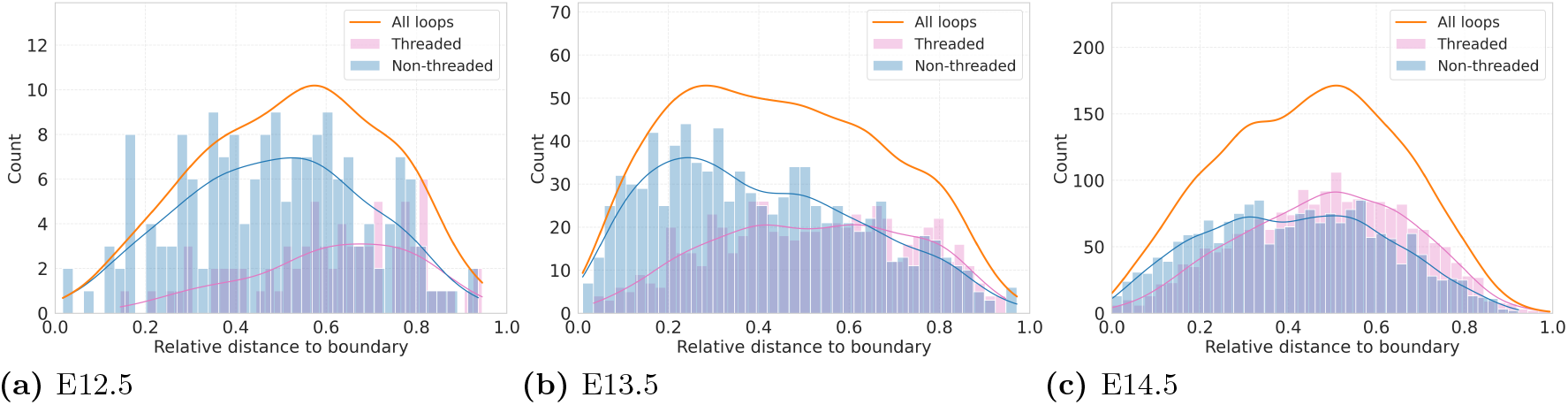
Distribution of the relative position of the death point related to each loop inside the pancreas in VAS. The orange line is the distribution built over all points, while pink corresponds to only the threaded loop via DUC, and blue to non-threaded. The plots are built over all the embryos separately for each day. The histograms do not have the same y-axis to emphasise the shape within each day.

## Discussion

In this work, we developed a computational pipeline to extract, pair, and classify loop structures in three pancreatic networks (neurons (NEU), ducts (DUC), and vasculature (VAS)) and applied chromatic persistence to quantify their spatial entanglement during development. Our analysis reveals several geometric trends that characterise the coordinated formation of these networks.

At first, we consider differential timing and spatial organisation in loop formation. Loop formation does not occur simultaneously across all networks. Loops in the ductal and vascular networks are already present at day E12.5, whereas loops in the neuronal network appear mostly at day E14.5, with only a few small loops at E13.5. This temporal offset suggests that the neuronal network establishes its loop architecture later in development, potentially after the ductal and vascular scaffolds are in place. Furthermore, the relative loop density in VAS remains stable over time, indicating that the vascular network does not remodel toward a tree-like topology. In contrast, DUC exhibits a decrease in relative loop density from day E13.5 to day E14.5, consistent with ductal remodelling and the emergence of a more hierarchical branching structure as described previously [1, 28].

We next examined the typical geometric scales associated with loop formation. The characteristic loop sizes are similar across networks, with peaks ranging from approximately 63 to 81 *µ*m. Notably, the loop size distribution in the vasculature shifts toward larger loops between day E12.5 and day E13.5, after which it stabilises. The similarity of distributions at days E13.5 and E14.5 suggests that the geometric characteristics of loop formation are largely established by day E13.5.

We next considered where loops are located within the organ and whether their geometry varies with position. We observe that loops in both the duct and vasculature are more densely concentrated toward the centre of the pancreas. However, we find no correlation between loop size and spatial position in NEU and VAS. No systematic correlation is observed for DUC either, yet it shows that the few largest loops are located closer to the centre of the organ.

Finally, we look at interactions between networks and quantify how their structures intertwine. A central contribution of this work is the quantification of inter-network entanglement through chromatic persistence. Our results reveal several interesting patterns with biological relevance:

- **The neural network is highly entangled with vasculature**. At day E14.5, approximately 84% of neuronal loops are threaded by vasculature, compared to 58% by ducts. This asymmetry, coupled with the temporal pattern where neuronal loops develops after vasculature, suggests that the vascular network provides the dominant spatial scaffold through which the neuronal network weaves, and not the other way around as seen in some other tissues. Interestingly, the high degree of threading seems less consistent with blood-vessel guided neuronal migration, as seen in the brain [29, 30], but may indicate a mechanism more consistent with the complex pull-push interaction with endothelial cells exhibited by motor neurons [31].
- **Ductal - vascular entanglement is mutual and increases over development**. Threading between ducts and vasculature is bidirectional, with approximately 40–50% of loops in each network threaded by the other at day 14.5. This proportion increases from day E12.5 onward, reflecting progressive spatial interweaving as both networks expand, in accordance with a close coordination of vasculature and pancreatic epithelium [5].
- **The neuronal network contributes minimally to threading of other networks**. While NEU loops are frequently threaded by DUC and VAS, the reverse is not true: only 5% of ductal loops and 15% of vascular loops are threaded by neurons. This asymmetry likely reflects the later developmental timing and smaller overall extent of the neuronal network. It would be relevant to analyse in later studies if the neuronal threading becomes more pronounced at later stages.
- **Larger loops are preferentially threaded**. Across all network pairs, threaded loops tend to be larger than non-threaded loops. This geometric bias may arise because larger loops present greater opportunities for spatial intersection with other structures. For ductal loops, this also aligns with our previous finding that larger ductal loops are more stable [1].
- **Threading is concentrated in the organ interior**. Loops located near the periphery of the pancreas tend to remain non-threaded, while loops closer to the centre are more frequently entangled. Since loop size does not correlate linearly with spatial position, this pattern reflects a genuine spatial organisation of network interactions rather than a size-dependent artefact. We find this result highly interesting in light of our previous finding that beta cells are preferentially found in large, centrally located loops, which provide a unique extracellular matrix (ECM) rich niche, and speculate that this ECM profile could be induced by threading vasculature [1].

On the computational side, we introduced a pipeline that combines discrete Morse skeletonisation with persistence-based loop filtering, reducing the number of candidate loops by approximately 90% while retaining topologically meaningful structures. The pairing algorithm provides a principled approach to matching persistence points with geometric representatives. Our application of chromatic persistence to classify loops as threaded or non-threaded offers a framework for quantifying spatial interactions between biological networks that may be applicable to other multi-component systems.

Several limitations should be noted. The sample sizes at each developmental stage (5, 3, and 3 embryos at days E12.5, E13.5, and E14.5, respectively) preclude formal statistical inference; our analysis is therefore descriptive rather than hypothesis-driven. For practical reasons we only analysed the dorsal part of the pancreas. It would be interesting to analyse if the ventral pancreas networks follow the same patterns. Methodologically, the chromatic approach cannot distinguish between topologically linked loops and loops that are merely in close spatial proximity—both configurations result in changes to the persistence diagram. However, for the purpose of characterising network entanglement, both scenarios indicate meaningful spatial interaction.

Several methodological considerations merit discussion. Geometric loop candidates obtained from skeleton processing and minimum cycle basis extraction do not always faithfully represent the underlying loops in the data, as loops in three dimensions can have complex shapes. Consequently, loop size, defined as path length along the skeleton, can exceed the actual size of the corresponding hole in the network.

Computational scale presents a further challenge: SEDT filtrations, persistent homology, and minimum cycle bases must be computed on subvolumes and subsequently merged with consistency checks on overlapping regions, whereas the pairing procedure is efficient enough to run on full samples.

Skeletonisation was performed on the raw anisotropic voxel grid (1.4 ×0.332 ×0.332 *µ*m), as interpolation to isotropic spacing would substantially increase memory without affecting the extracted loop topology. Loops are rescaled to physical coordinates before pairing. In contrast, SEDT and chromatic persistence computations are metric-dependent and explicitly account for voxel spacing throughout.

Finally, we note that the biological data exhibit considerable variability between embryos; we therefore present multiple representative samples and provide per-embryo plots in the Supporting Information.

## Acknowledgments

The authors would like to thank Yossi Bokor Bleile, Elodie Maignant, and Vanessa Robins for valuable discussions and insightful comments that contributed to this work. We acknowledge the Danstem Imaging platform at University of Copenhagen for use of the confocal microscope and the Microscope Core Facility at Department of Science and Environment, Roskilde University for use of image analysis software. P Nyeng is supported by a research grant (VIL69255) from Villum Fonden.

## References

1. Jackson AL, Heilmann S, Agerskov R, Ebeid C, Krivokapic JM, Romero Herrera JA, et al. Real-time imaging reveals new mechanisms for pancreatic ductal establishment and remodeling. Journal of Cell Biology. 2026;225(3):e202409022.

2. Bankaitis ED, Bechard ME, Wright CV. Feedback control of growth, differentiation, and morphogenesis of pancreatic endocrine progenitors in an epithelial plexus niche. Genes & development. 2015;29(20):2203–2216.

3. Agerskov RH, Nyeng P. Innervation of the pancreas in development and disease. Development. 2024;151(2):dev202254.

4. Alvarsson A, Jimenez-Gonzalez M, Li R, Rosselot C, Tzavaras N, Wu Z, et al. A 3D atlas of the dynamic and regional variation of pancreatic innervation in diabetes. Science Advances. 2020;6(41):eaaz9124.

5. Cleaver O, Dor Y. Vascular instruction of pancreas development. Development. 2012;139(16):2833–2843.

6. Pierreux CE, Cordi S, Hick AC, Achouri Y, De Almodovar CR, Prévot PP, et al. Epithelial: Endothelial cross-talk regulates exocrine differentiation in developing pancreas. Developmental biology. 2010;347(1):216–227.

7. Reinert RB, Cai Q, Hong JY, Plank JL, Aamodt K, Prasad N, et al. Vascular endothelial growth factor coordinates islet innervation via vascular scaffolding. Development. 2014;141(7):1480–1491.

8. Edelsbrunner, Letscher, Zomorodian. Topological persistence and simplification. Discrete & Computational Geometry. 2002;28:511–533.

9. Frosini P. A distance for similarity classes of submanifolds of a Euclidean space. Bulletin of the Australian Mathematical Society. 1990;42(3):407–415.

10. Robins V. Towards computing homology from finite approximations. In: Topology Proceedings. vol. 24; 1999. p. 503–532.

11. Cohen-Steiner D, Edelsbrunner H, Harer J, Morozov D. Persistent homology for kernels, images, and cokernels. In: Proceedings of the twentieth annual ACM-SIAM symposium on Discrete algorithms. SIAM; 2009. p. 1011–1020.

12. di Montesano SC, Draganov O, Edelsbrunner H, Saghafian M. Chromatic Topological Data Analysis. arXiv preprint arXiv:240604102. 2024;.

13. Herring A, Robins V, Sheppard A. Topological persistence for relating microstructure and capillary fluid trapping in sandstones. Water Resources Research. 2019;55(1):555–573.

14. Raichenko V, Rosenthal N, Eder M, Evans ME. Cocoon microstructures through the lens of topological persistence. Journal of the Royal Society Interface. 2024;21(220):20240218.

15. Lee Y, Barthel SD, Dlotko P, Moosavi SM, Hess K, Smit B. Quantifying similarity of pore-geometry in nanoporous materials. Nature communications. 2017;8(1):1–8.

16. Christensen R, Bokor Bleile Y, Sørensen SS, Biscio CA, Fajstrup L, Smedskjaer MM. Medium-range order structure controls thermal stability of pores in zeolitic imidazolate frameworks. The Journal of Physical Chemistry Letters. 2023;14(33):7469–7476.

17. Bleile YB, Yadav P, Koehl P, Rehfeldt F. Persistence diagrams as morphological signatures of cells: A method to measure and compare cells within a population. PLOS Computational Biology. 2026;22(1):e1013890.

18. Stolz BJ, Dhesi J, Bull JA, Harrington HA, Byrne HM, Yoon IH. Relational persistent homology for multispecies data with application to the tumor microenvironment. Bulletin of Mathematical Biology. 2024;86(11):128.

19. Stolz B, Kaeppler J, Markelc B, et al. Multiscale topology characterizes dynamic tumor vascular networks. Sci Adv 8 (23): eabm2456; 2022.

20. Palla G, Fischer DS, Regev A, Theis FJ. Spatial components of molecular tissue biology. Nature Biotechnology. 2022;40(3):308–318.

21. Landuzzi F, Nakamura T, Michieletto D, Sakaue T. Persistence homology of entangled rings. Physical Review Research. 2020;2(3):033529.

22. Heilmann S, Semb H, Nyeng P. Quantifying spatial position in a branched structure in immunostained mouse tissue sections. STAR protocols. 2021;2(4):100806.

23. Delgado-Friedrichs O, Robins V, Sheppard A. Skeletonization and partitioning of digital images using discrete morse theory. IEEE transactions on pattern analysis and machine intelligence. 2014;37(3):654–666.

24. Robins V, Wood PJ, Sheppard AP. Theory and algorithms for constructing discrete Morse complexes from grayscale digital images. IEEE Transactions on pattern analysis and machine intelligence. 2011;33(8):1646–1658.

25. Lee TC, Kashyap RL, Chu CN. Building skeleton models via 3-D medial surface axis thinning algorithms. CVGIP: graphical models and image processing. 1994;56(6):462–478.

26. Kavitha T, Mehlhorn K, Michail D, Paluch KE. An algorithm for minimum cycle basis of graphs. Algorithmica. 2008;52(3):333–349.

27. Hagberg A, Swart PJ, Schult DA. Exploring network structure, dynamics, and function using NetworkX. Los Alamos National Laboratory (LANL), Los Alamos, NM (United States); 2008.

28. Kesavan G, Sand FW, Greiner TU, Johansson JK, Kobberup S, Wu X, et al. Cdc42-mediated tubulogenesis controls cell specification. Cell. 2009;139(4):791–801.

29. Fujioka T, Kaneko N, Sawamoto K. Blood vessels as a scaffold for neuronal migration. Neurochemistry international. 2019;126:69–73.

30. Ogino T, Saito A, Sawada M, Takemura S, Hara Y, Yoshimura K, et al. Neuronal migration depends on blood flow in the adult mammalian brain. Elife. 2025;13:RP99502.

31. Martins LF, Brambilla I, Motta A, de Pretis S, Bhat GP, Badaloni A, et al. Motor neurons use push-pull signals to direct vascular remodeling critical for their connectivity. Neuron. 2022;110(24):4090–4107.

