## Supplementary Material for "Topological Analysis of Multi-Network Threading in the Pancreas"

Vira Raichenko[1], Rebin Maaruf[2], Pia Nyeng[2], Myfanwy Evans [1*],
Associate Professor, Department of Science and Environment Molecular and Medical Biology Centre for Mathematical Modelling - Human Health and Disease

**1** Institute for Mathematics, University of Potsdam, Potsdam, Brandenburg, Germany
**2** Department of Science and Environment Molecular and Medical Biology Centre for Mathematical Modelling - Human Health and Disease, Roskilde University, Roskilde, Denmark

*

### 1 Supporting Information

This Supporting Information is organised into two parts. The first part presents per-embryo plots of the analyses reported in the main text. Since the main text aggregates data across embryos at each developmental stage, these individual plots allow the reader to assess variability between samples. They include loop size distributions, loop size versus distance to the organ boundary, spatial density maps of loops, size distributions of threaded loops, and the spatial distribution of threaded versus non-threaded loops relative to the organ boundary, all shown separately for each embryo and network type. The second part provides a detailed description of the persistence pair to loop cycle matching algorithm. This includes a visual walkthrough of the pairing procedure on a small example, the formal algorithmic description with pseudocode for candidate harvesting, loop scoring, primary selection, and collision resolution, and a table summarising all algorithm parameters used in the analysis.

#### 1.1 Size and spatial distribution plots per embryo

In the paper "Topological Analysis of Multi-Network Threading in Pancreas", we presented an analysis of neuronal, ductal, and vascular networks. The dataset consisted of five embryos at day E12.5 of pancreatic development, three embryos at day E13.5, and three embryos at day E14.5. In that work, most plots were generated by combining data from all embryos at the same developmental stage. However, as individual-embryo variability may be of interest, we provide here the corresponding plots for each embryo separately.

To illustrate the variability across embryos, we present the size distributions of loops in NEU (Figure 1), DUC (Figure 4), and VAS (Figure 5). Each figure shows the loop size distribution for individual embryos in the dataset, including data from all developmental days. The legend reports the total number of loops in each histogram. The axes are kept uniform across figures; as a result, the histogram shapes for day E12.5 are difficult to discern. To address this, we include an enlarged inset highlighting this region.

We plot loop size against the positions of the corresponding death points, which provide a better representation of loop centers than the center of mass, as many loops are non-convex. The corresponding scatter plots are shown for NEU (Figure 6), DUC (Figure 7), and VAS (Figure 8).

Finally, the relative positions of loops within the pancreas are shown as histograms for the two loop classes, threaded and non-threaded. The corresponding figures present DUC loops threaded via VAS (Figure 14) and VAS loops threaded via DUC (Figure 15).

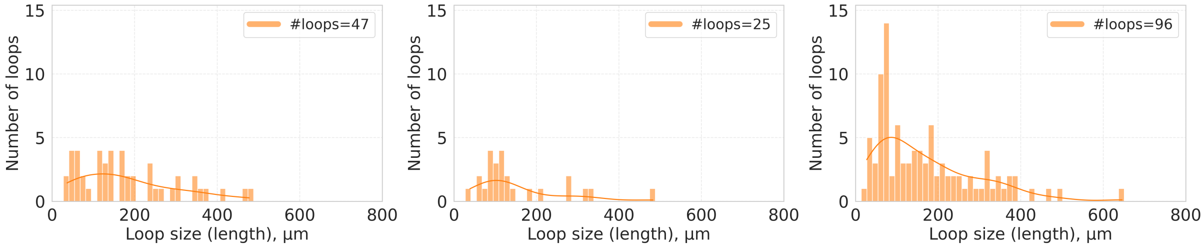


**(a)** **(b)** **(c)**

**Fig 1.** Loop size distributions for NEU for each embryo at day E14.5. The legend provides the total number of loops.

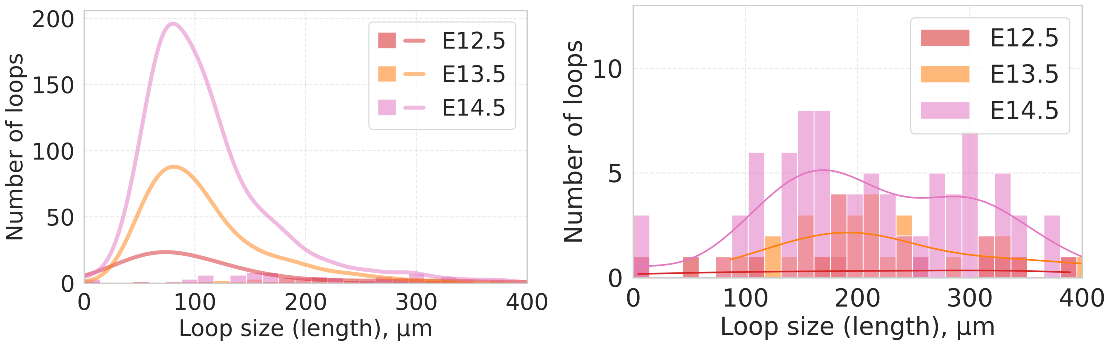


**(a)** C2 via C3 **(b)** C3 via C2

**Fig 2.** Distribution of threaded loop size in DUC via NEU. (a) The histogram displays the distribution of threaded loops on each day. The approximation of the distribution plotted as lines corresponds to the distributions of the sizes of all loops for each day, respectively. The colors of the lines correspond to the colors of the histograms for each day accordingly. (b) The same histogram as in (a) but on a different y-axis scale for better display of the shapes of the histograms with respect to each other. In both cases, each plot is built over all embryos for each day.

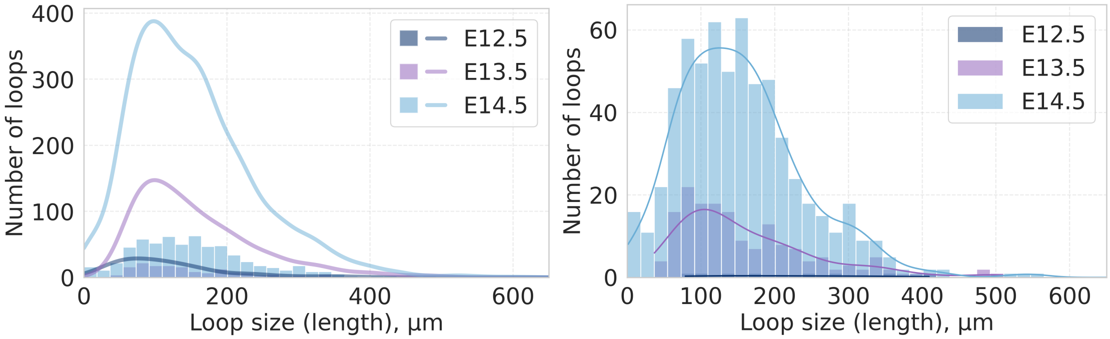


**(a)** C2 via C3 **(b)** C3 via C2

**Fig 3.** Distribution of threaded loop size in VAS via NEU. (a) The histogram displays the distribution of threaded loops on each day. The approximation of the distribution plotted as lines corresponds to the distributions of the sizes of all loops for each day, respectively. The colors of the lines correspond to the colors of the histograms for each day accordingly. (b) The same histogram as in (a) but on a different y-axis scale for better display of the shapes of the histograms with respect to each other. In both cases, each plot is built over all embryos for each day.

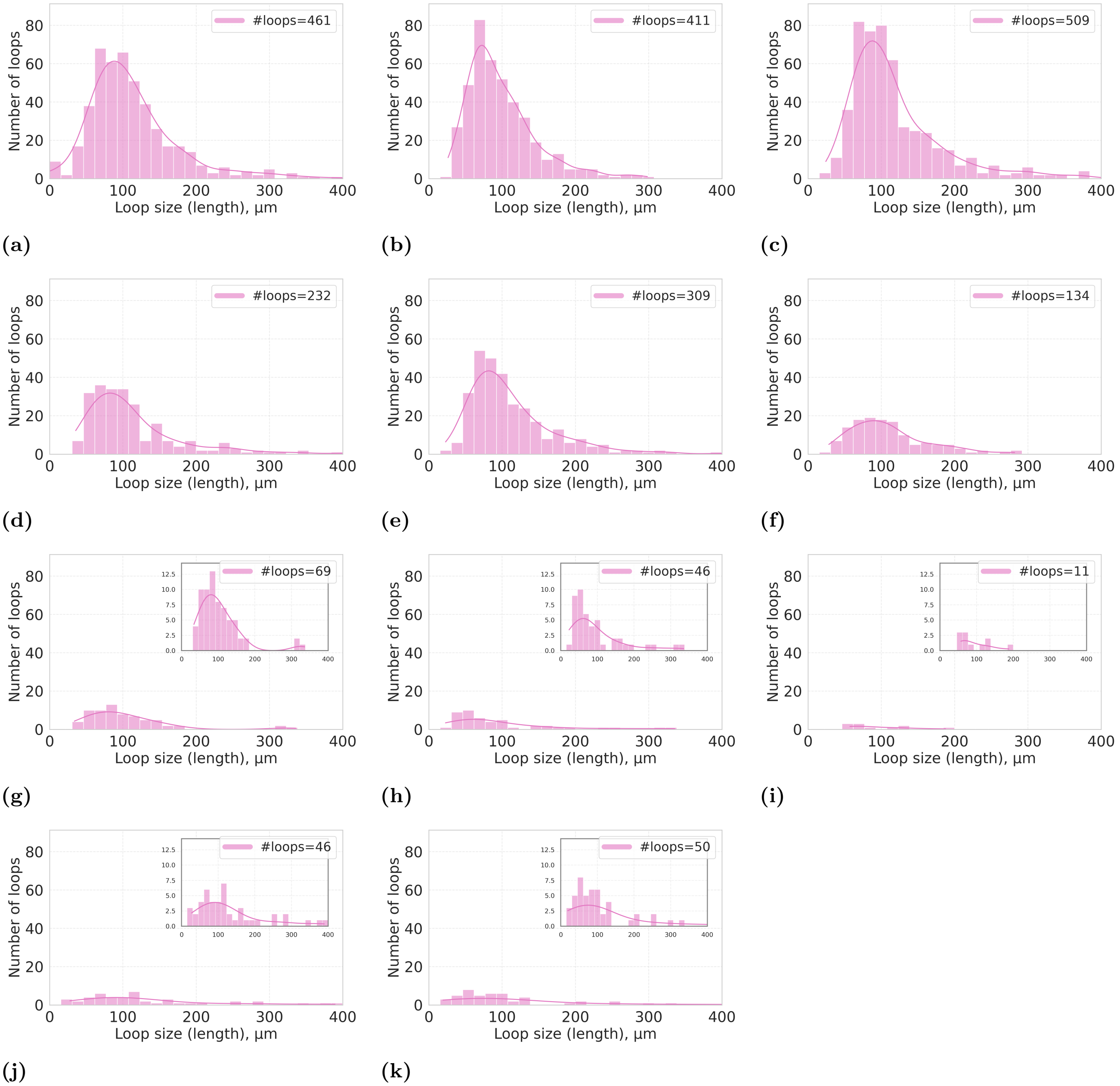


**Fig 4.** Loop size distributions for DUC for each embryo: (a)–(c) day E14.5, (d)–(f) day E13.5, and (g)–(k) day E12.5. The legend provides the total number of loops.

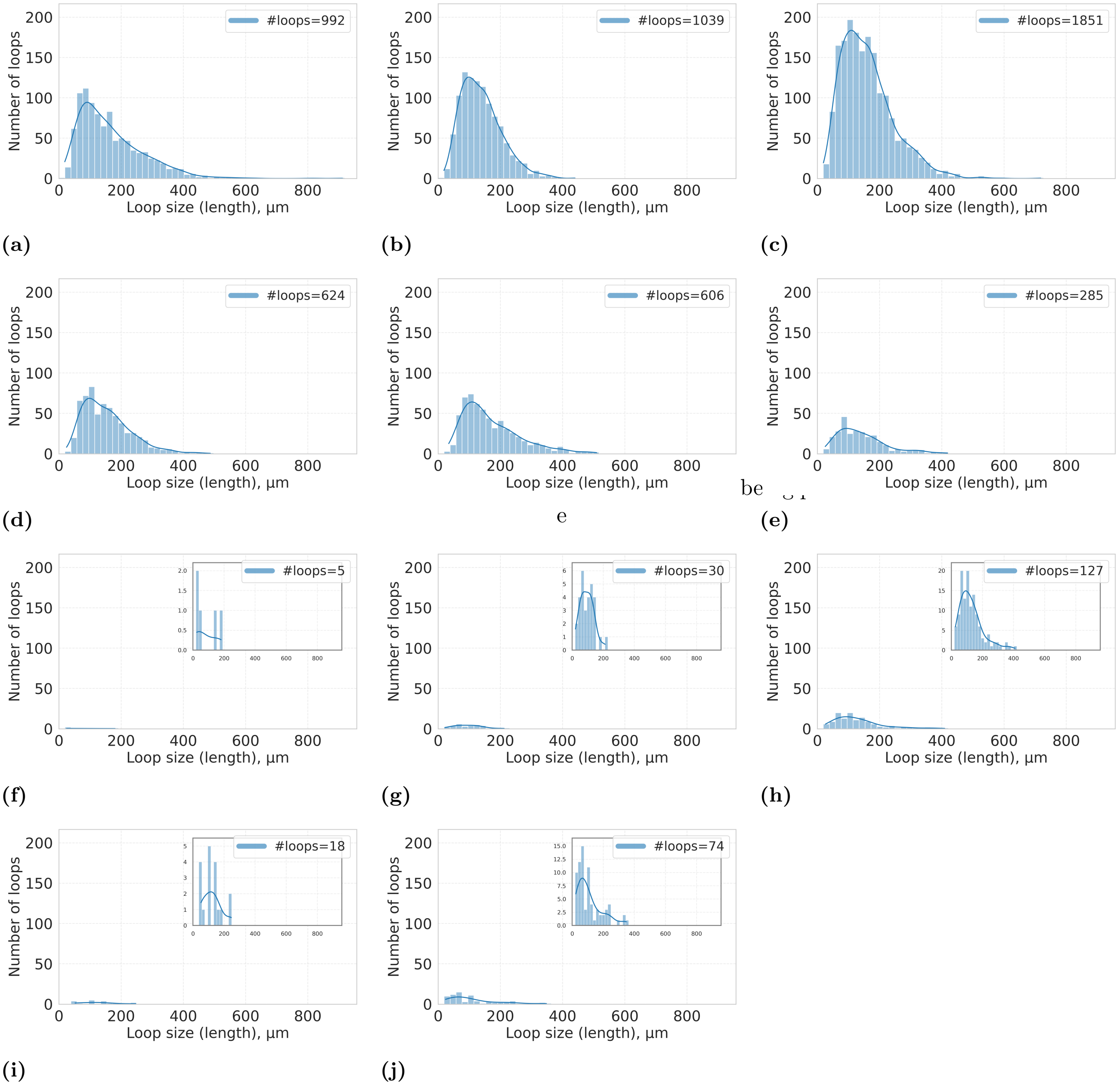


**Fig 5.** Loop size distributions for VAS for each embryo: (a)–(c) day E14.5, (d)–(f) day E13.5, and (g)–(k) day E12.5. The legend provides the total number of loops.

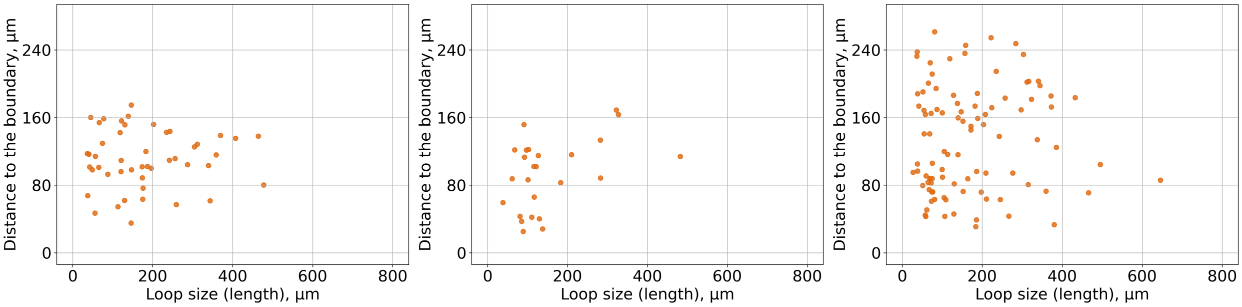


**(a)** **(b)** **(c)**

**Fig 6.** Loops size vs distance to sample per embryo in NEU day E14.5.

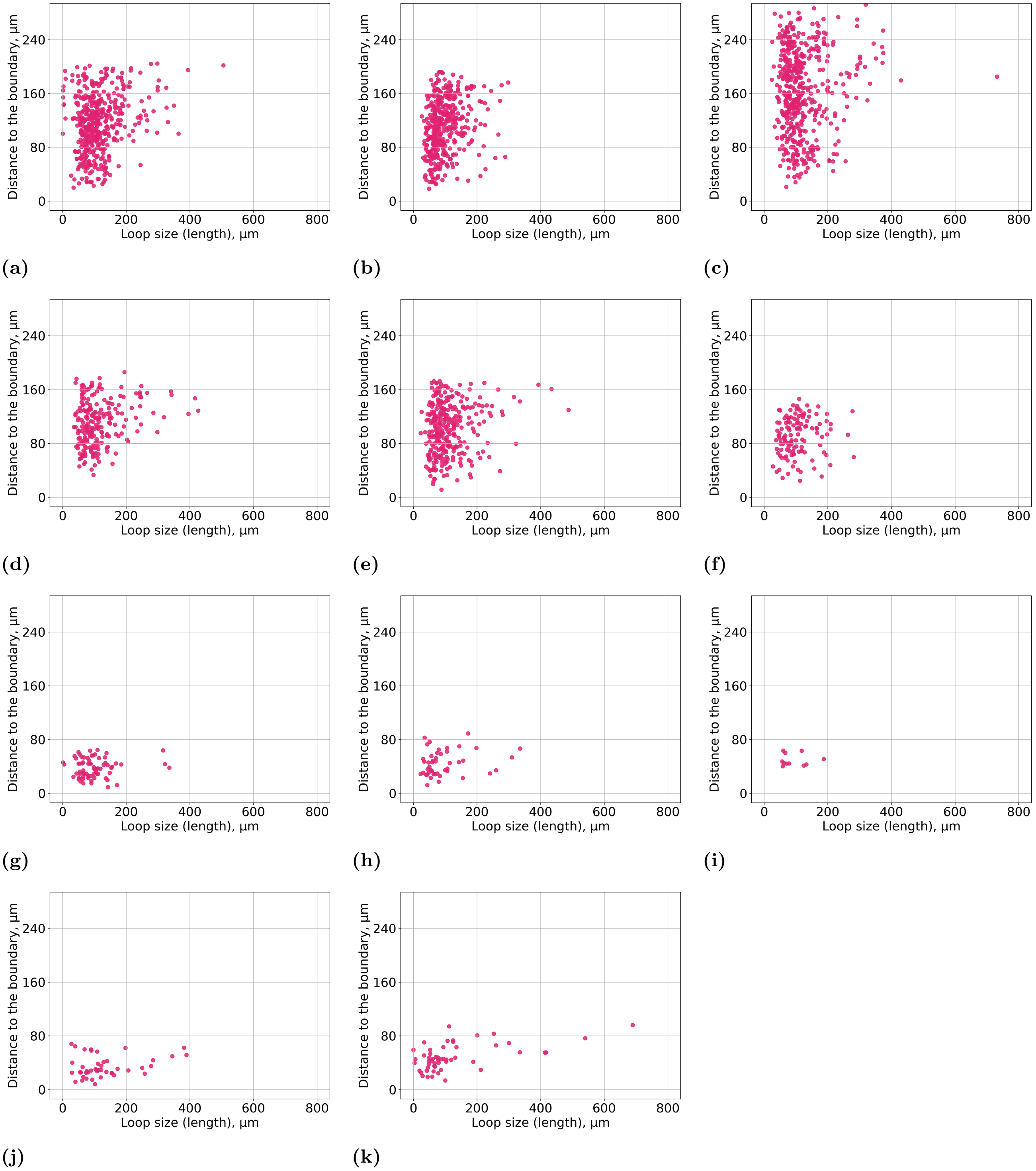


**Fig 7.** Loops size vs distance to sample per embryo in DUC. (a)-(c) day E14.5, (d)-(f) day E13.5 and (g)-(k) day E12.5.

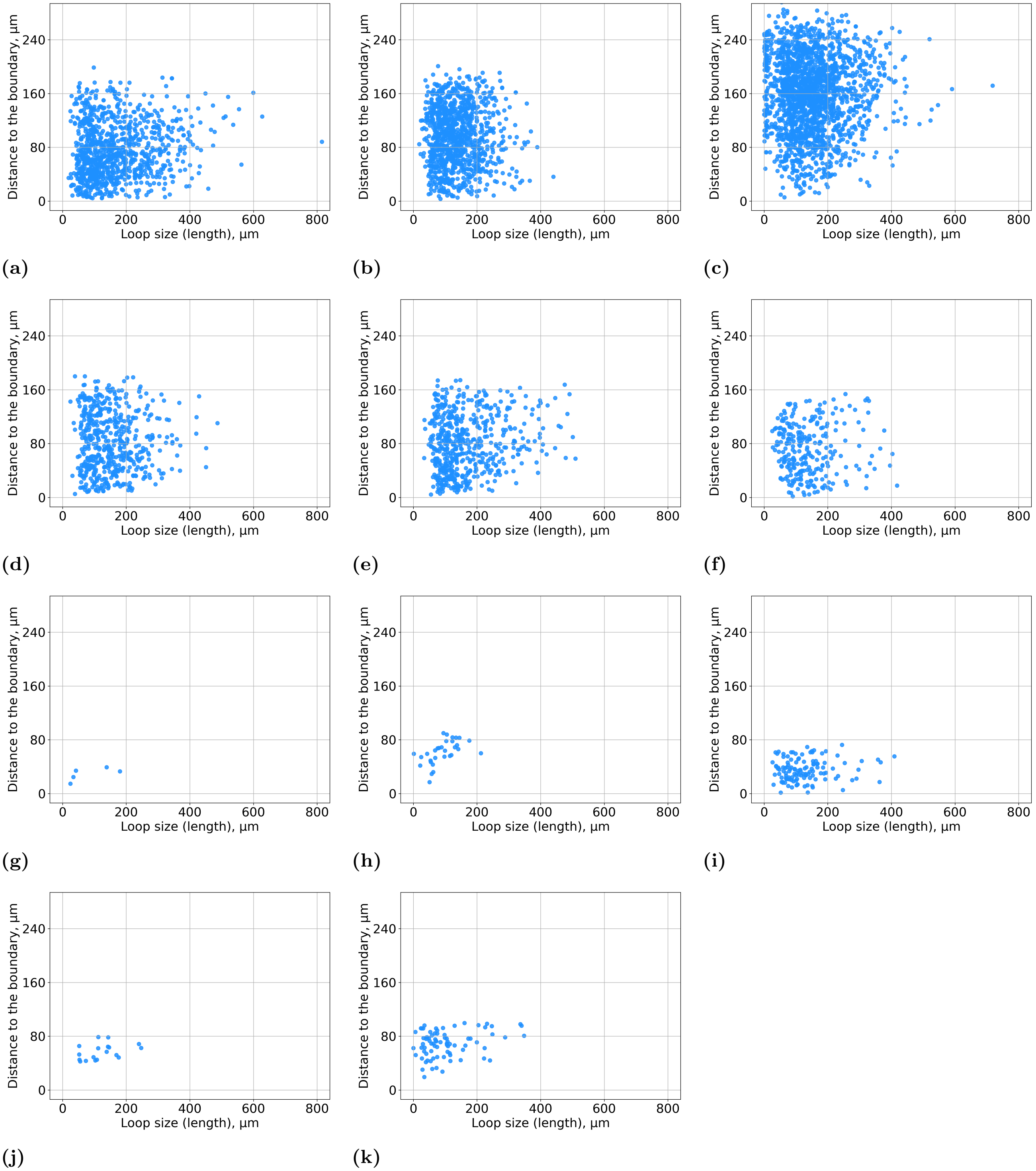


**Fig 8.** Loops size vs distance to sample per embryo in VAS. (a)-(c) day E14.5, (d)-(f) day E13.5 and (g)-(k) day E12.5.

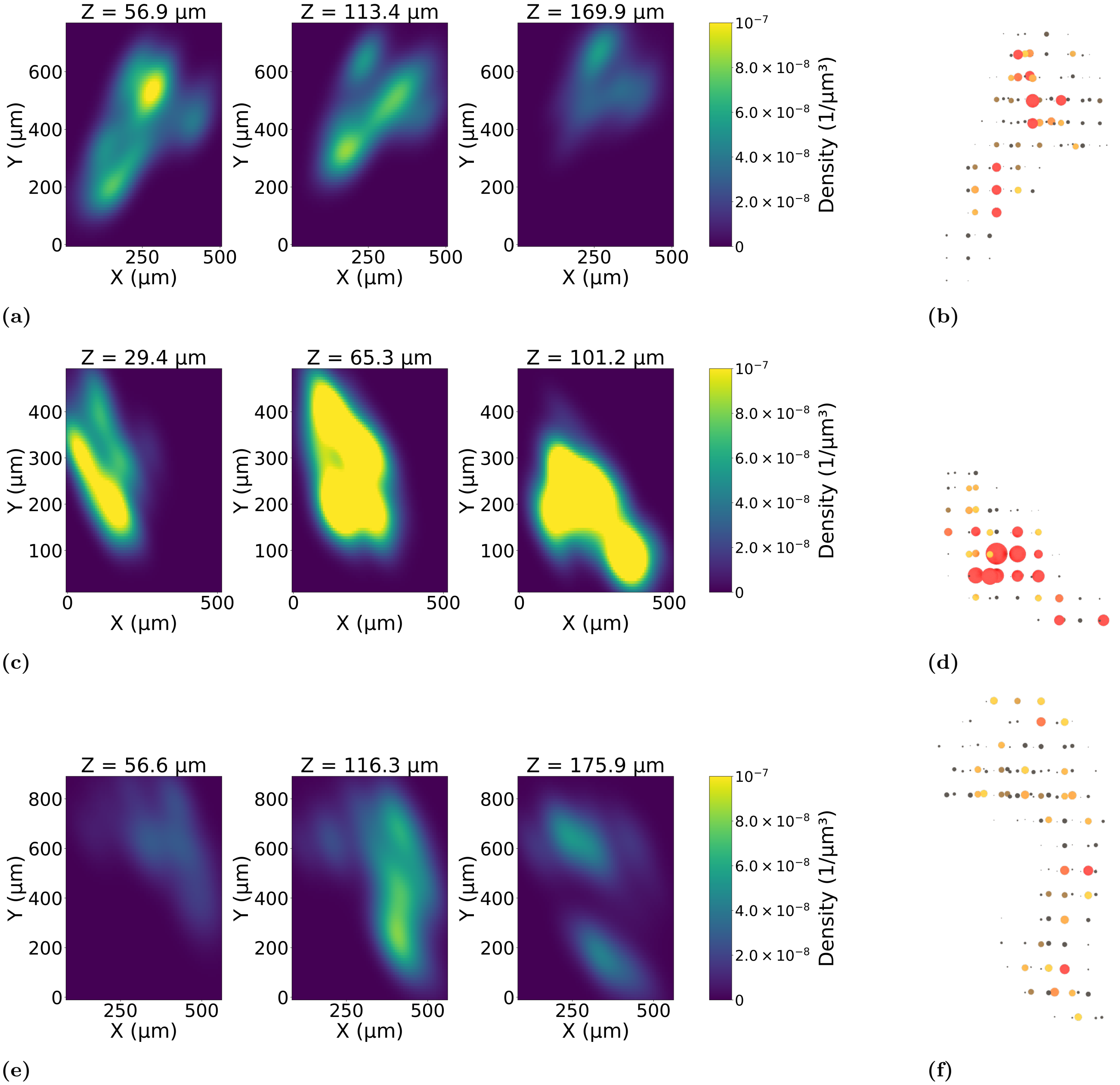


**Fig 9.** The density of loop distributions in DUC on the day E14.5. Panels (a), (c), and (e) correspond to different samples. In each row, the middle slice is taken at the centre of the volume along the shortest axis ($z$), while the left and right slices are taken at $\pm 20$ slices. The intensity scale is kept constant across all samples to demonstrate variability between samples. Panels (b), (d), and (f) show schematic 3D histogram representations: bins correspond to cubes of size 60 per side, and the radius of each sphere represents the number of loop centres (death points) within the corresponding cube. Colouring highlights regions with a high concentration of loops, with red indicating the highest densities.

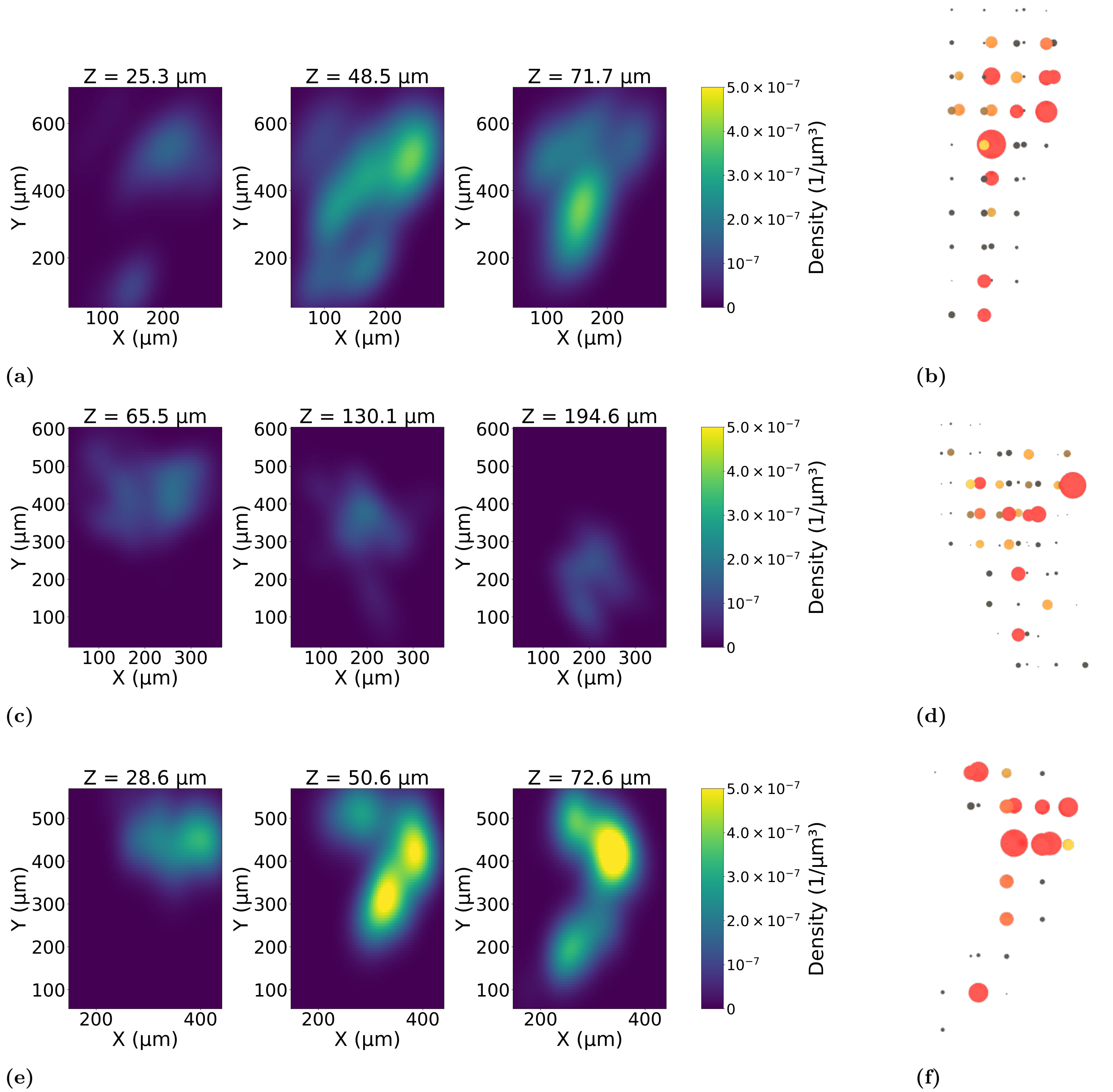


**Fig 10.** The density of loop distributions in DUC on the day E13.5. Panels (a), (c), and (e) correspond to different samples. In each row, the middle slice is taken at the centre of the volume along the shortest axis ($z$), while the left and right slices are taken at ±20 slices. The intensity scale is kept constant across all samples to demonstrate variability between samples. Panels (b), (d), and (f) show schematic 3D histogram representations: bins correspond to cubes of size 60 per side, and the radius of each sphere represents the number of loop centres (death points) within the corresponding cube. Colouring highlights regions with a high concentration of loops, with red indicating the highest densities.

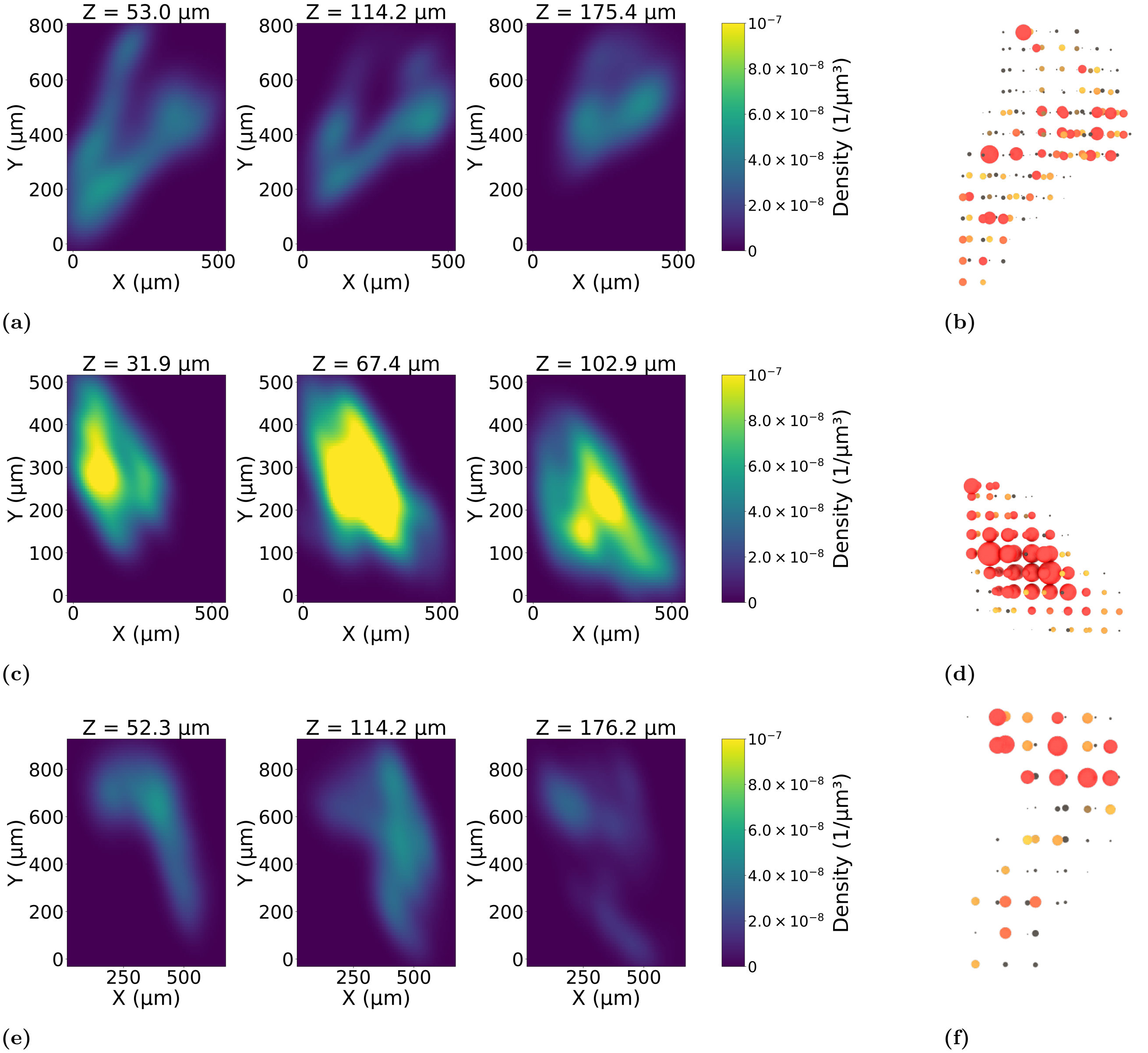


**Fig 11.** The density of loop distributions in VAS on the day E14.5. Panels (a), (c), and (e) correspond to different samples. In each row, the middle slice is taken at the centre of the volume along the shortest axis ($z$), while the left and right slices are taken at $\pm 20$ slices. The intensity scale is kept constant across all samples to demonstrate variability between samples. Panels (b), (d), and (f) show schematic 3D histogram representations: bins correspond to cubes of size 60 per side, and the radius of each sphere represents the number of loop centres (death points) within the corresponding cube. Colouring highlights regions with a high concentration of loops, with red indicating the highest densities.

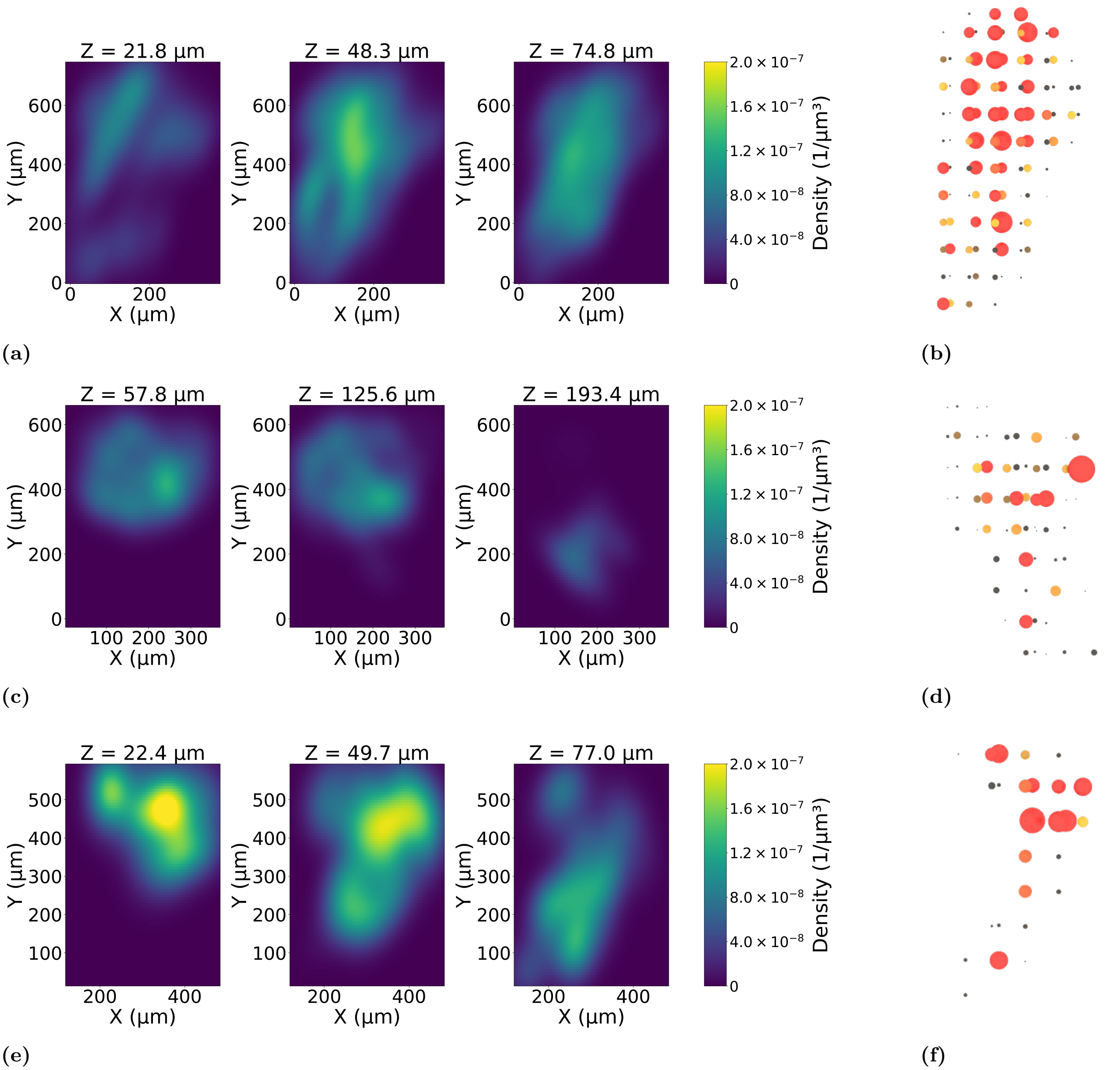


**Fig 12.** The density of loop distributions in VAS on the day E13.5. Panels (a), (c), and (e) correspond to different samples. In each row, the middle slice is taken at the centre of the volume along the shortest axis ($z$), while the left and right slices are taken at $\pm 20$ slices. The intensity scale is kept constant across all samples to demonstrate variability between samples. Panels (b), (d), and (f) show schematic 3D histogram representations: bins correspond to cubes of size 60 per side, and the radius of each sphere represents the number of loop centres (death points) within the corresponding cube. Colouring highlights regions with a high concentration of loops, with red indicating the highest densities.

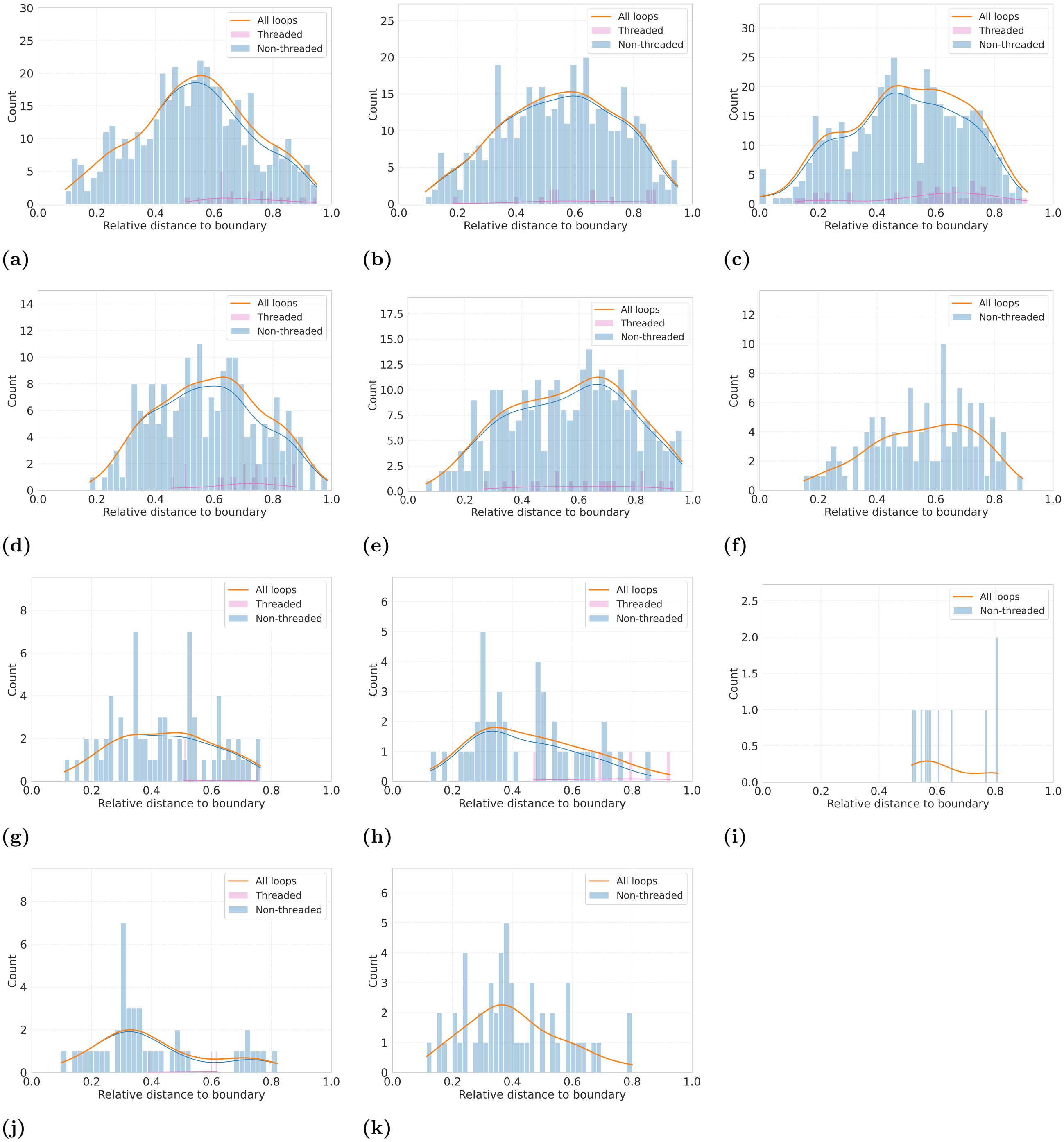


**Fig 13.** Threaded and non-threaded loops vs distance to sample per embryo in DUC via NEU. (a)-(c) day E14.5, (d)-(f) day E13.5 and (g)-(k) day 12.5. If "Threaded" label is missing, it means no loops of C2 got threaded via C1.

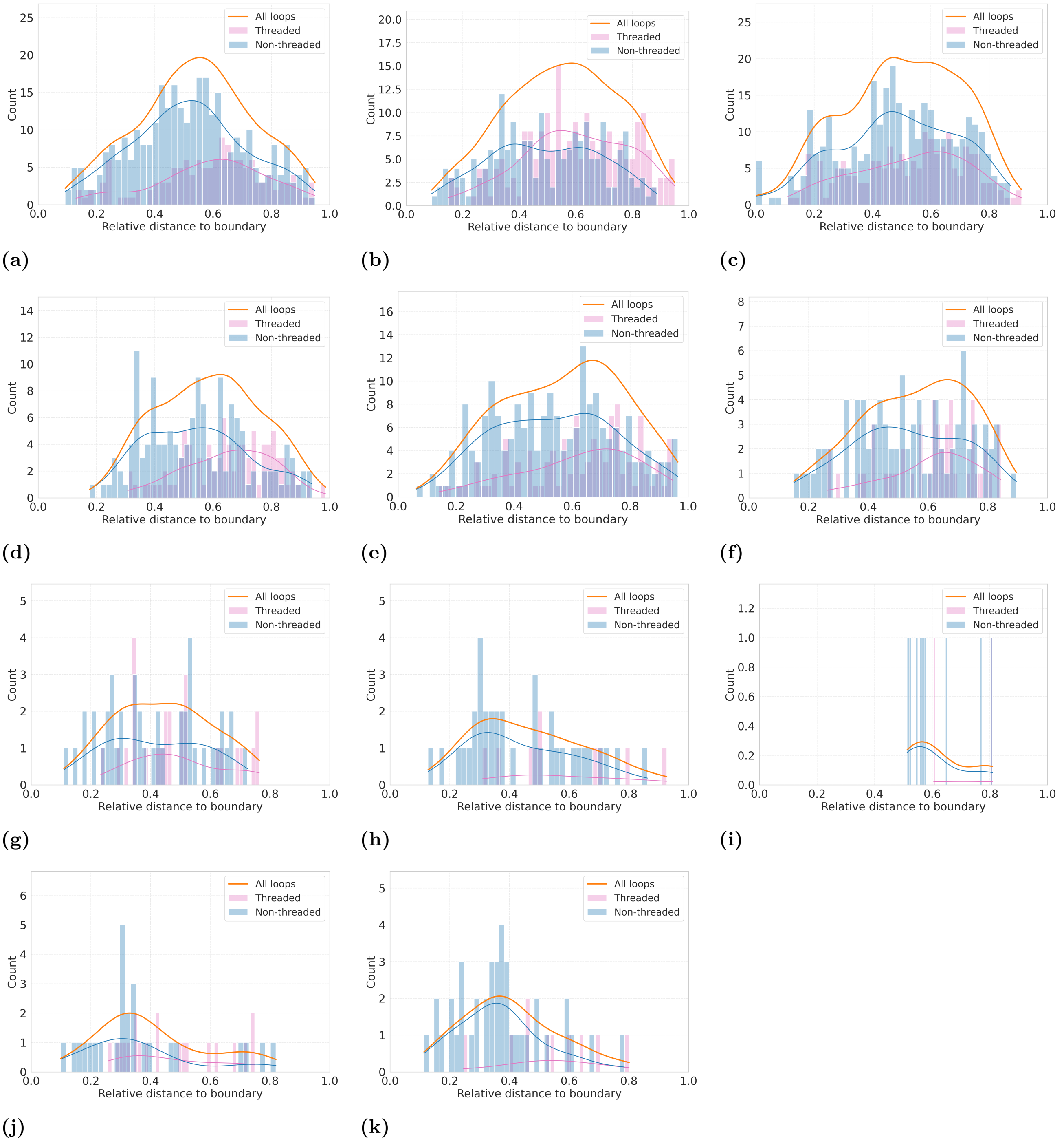


**Fig 14.** Threaded and non-threaded loops vs distance to sample per embryo in DUC via VAS. (a)-(c) day E14.5, (d)-(f) day E13.5 and (g)-(k) day E12.5.

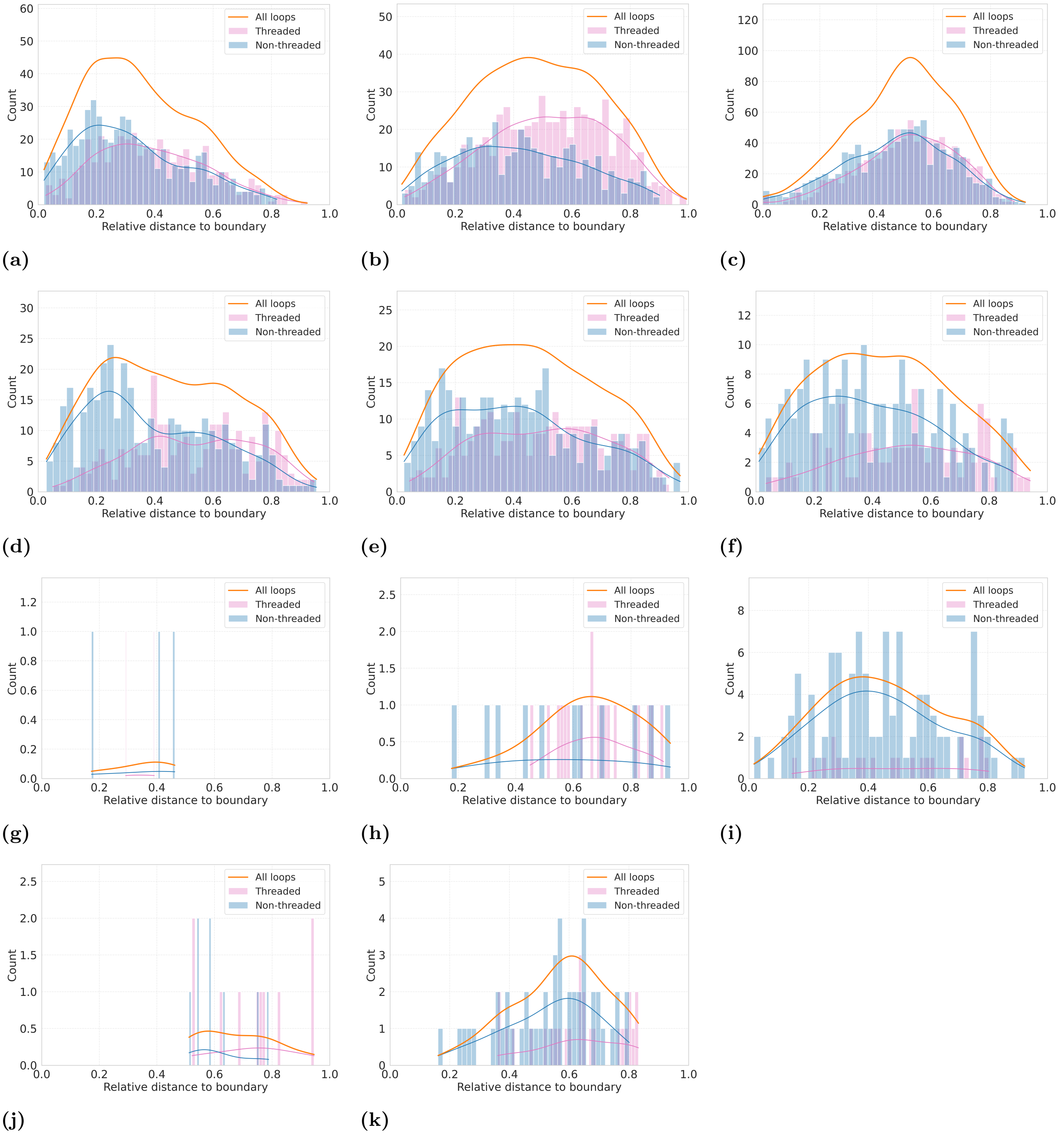


**Fig 15.** Threaded and non-threaded loops vs distance to sample per Embryo in VAS via DUC. (a)-(c) day E14.5, (d)-(f) day E13.5 and (g)-(k) day E12.5.

#### 1.2 Pairing algorithm

As an essential step of our pipeline, we pair the geometric loops extracted via a minimum cycle basis (i.e., a cycle basis that minimises the total weight/length of all cycles) with points in the persistence diagrams. To provide intuition for the choices made by the pairing algorithm, we give a more detailed overview how it makes decisions on a simple example of loops taken from the data. A visual example of a sample with loops and their corresponding (birth, death) pairs that we consider is in Figure 16. We consider 3D loops from NEU network, and we choose the combination of the loops that are relatively straightforward to pair; however, loops in DUC and VAS can be much more complicated.

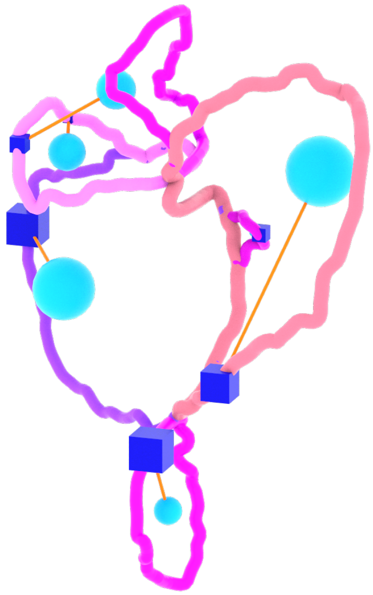

**Fig 16.** A set of geometric loops and a set of birth(dark blue cubes) with corresponding death(light blue spheres) that require a matching The orange connections are between (birth, death) pairs. The pairing algorithm aims to match for each (birth, death) the best possible geometric loop.

##### 1.2.1 Persistence Pair to Loop Cycle Matching Algorithm

Given a persistence diagram $\mathcal{D} = \{(b_i, d_i)\}_{i=1}^{n}$ where $b_i \in \mathbb{R}^3$ denotes a birth point and $d_i \in \mathbb{R}^3$ denotes the corresponding death point, and a minimum cycle basis $\mathcal{L} = \{L_1, L_2, \ldots, L_m\}$ of the network graph, we seek an injective assignment $\phi : \mathcal{D} \to \mathcal{L} \cup \{\perp\}$ where $\perp$ denotes "unpaired."

##### 1.2.2 Candidate Harvesting

For each persistence pair $(b_i, d_i)$ with death scalar $\delta_i$, we construct a candidate set $\mathcal{C}_i \subseteq \mathcal{L}$ by combining two spatial queries: a $k$-nearest neighbor search around the birth point $b_i$ augmented with iterative radius expansion if needed, and a radius search around the death point $d_i$ with radius proportional to $|\delta_i|+$ offset. The union of loops found by both queries forms the final candidate set $\mathcal{C}_i$.

Figure 17 shows the highest-ranked candidate loops for three representative (birth, death) pairs. In the typical case Figures 17a–17b), the birth point lies on a loop that coincides with the eventual best match (Figure 19). In the third case (Figure 17c), however, the birth lies on a different loop than the one selected as the best match. This illustrates that the birth point can be far from the best-matching loop, which motivates the two-step candidate harvesting: the $k$-nearest neighbor search around the birth guarantees that loops passing through the birth are considered, while the radius search around the death ensures that loops enclosing the death point are included. Both pools are needed because neither alone reliably contains the correct match.

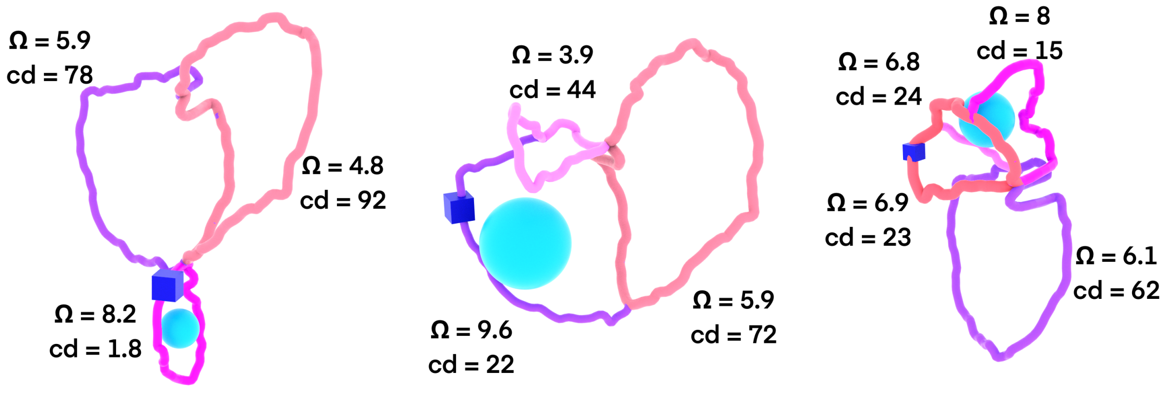


**(a)** **(b)** **(c)**

**Fig 17.** Top-ranked candidate loops for three representative (birth, death) pairs, i.e, candidates from the set $\mathcal{C}_i$. Each loop is annotated with the spherical arc length $\Omega$ (rad) and the centre distance cd from the loop barycenter to the death point. In each subfigure, the loop with the highest $\Omega$ and smallest cd corresponds to the visually best-matching candidate. Birth and death points are indicated by squares.

##### 1.2.3 Loop Scoring

For each candidate loop $L_j \in \mathcal{C}_i$, we compute three metrics. The **spherical arc length** measures the total geodesic arc length of the loop projected onto the unit sphere centred at the death point:

$$\Omega(L_j, d_i) = 2 \sum_{k=1}^{|L_j|} \arctan\left(\frac{\|\hat{v}_k \times \hat{v}_{k+1}\|}{1 + \hat{v}_k \cdot \hat{v}_{k+1}}\right) \tag{1}$$

where $\hat{v}_k = (v_k - d_i)/\|v_k - d_i\|$ and $v_k$ are vertices of $L_j$. The **center distance** is $\mathrm{cd}(L_j, d_i) = \|c_j - d_i\|$ where $c_j$ is the loop barycenter. The **minimum death distance** is $\mathrm{md}(L_j, d_i) = \min_{v \in L_j} \|v - d_i\|$.

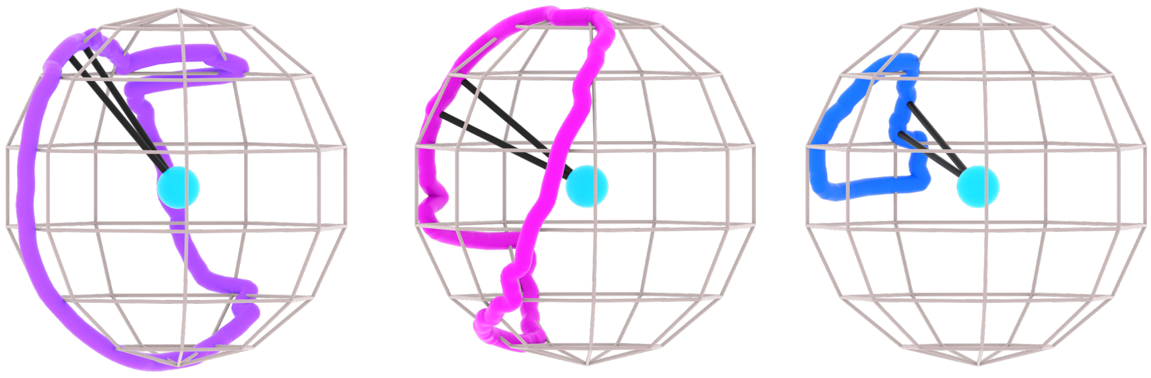

**(a)** $\Omega = 9.6$ **(b)** $\Omega = 9$ **(c)** $\Omega = 3.5$

**Fig 18.** Spherical arc length demonstration for different loops. Each loop is projected onto the unit sphere centred at the death point, and $\Omega$ is the total geodesic arc length of the resulting spherical curve. The black straight lines correspond to $\hat{v}_k$ and $\hat{v}_{k+1}$. A loop that wraps closely around the death point traces a long path on the sphere (large $\Omega$), whereas a loop far from the death point produces a short path (small $\Omega$). Since $\Omega$ measures perimeter rather than enclosed area, it can naturally exceed $2\pi$.

**Remark.** We choose arc length over, for example, solid angles since it is more robust than true solid angle for the given data, and because 3D loops are non-convex and their spherical projections can self-intersect. True solid angle would suffer from cancellation (overlapping regions subtracting) and orientation ambiguity, while arc length stays well-defined and monotonic.

Figures 18a-18b illustrate loops with high spherical arc length $\Omega$, where the corresponding death point lies approximately inside the loop. In each case, the loop and death point are translated so that the death point coincides with the origin, and the loop vertices are then projected onto the unit sphere. The last panel (Figure 18c) shows an example in which the death point lies outside the loop, resulting in a low $\Omega$ score.

##### 1.2.4 Primary Selection

We first apply a hard constraint, retaining only candidates satisfying $\mathrm{md}(L_j, d_i) \leq |\delta_i| + \epsilon$ to form the eligible set $\mathcal{E}_i$. Let $\mathcal{Q}_i = \{L_j \in \mathcal{E}_i : \Omega(L_j, d_i) \geq \Omega_{\text{floor}}\}$ denote the $\Omega$-qualified subset. The selection rule is:

$$L^* = \begin{cases} \arg\min_{L_j \in \mathcal{Q}_i} \mathrm{cd}(L_j, d_i) & \text{if } \mathcal{Q}_i \neq \emptyset \\ \arg\max_{L_j \in \mathcal{E}_i} \Omega(L_j, d_i) & \text{otherwise} \end{cases} \tag{2}$$

That is, if any candidate has spherical arc length above the threshold $\Omega_{\text{floor}}$, we select the one closest to the death point; otherwise we fall back to the candidate with maximum spherical arc length, see Figure 17. We also store a ranked preference list $\mathcal{P}_i$ of all eligible candidates for use in collision resolution.

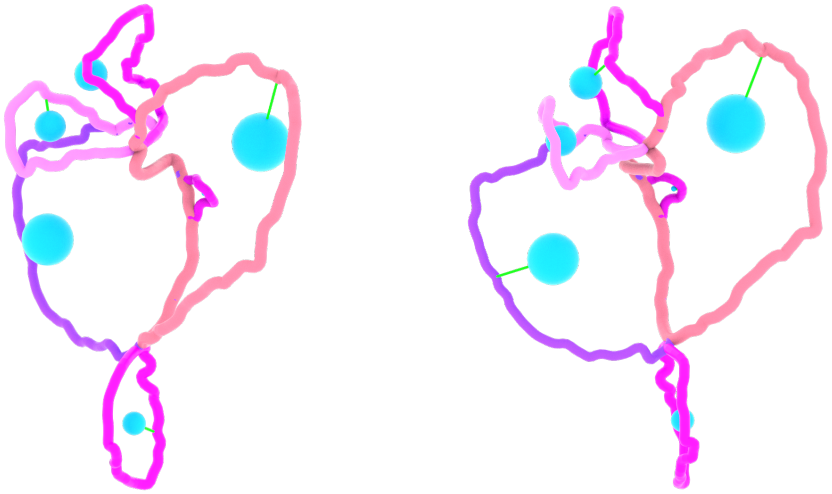

**(a)** **(b)**

**Fig 19.** Demonstration of the pairing. The green connections between the loops and the spheres correspond to the matching between the corresponding death and the loop. We demonstrate the same set of loops from different angles (a) and (b).

##### 1.2.5 Collision Resolution

The initial assignment may be non-injective, with multiple births assigned to the same loop. We resolve such collisions by identifying a *keeper* for each contested loop and attempting to reassign the remaining births to alternative loops.

For each loop $L_j$ claimed by multiple births $B_j$, we first determine which births can feasibly move to an alternative. A birth $i$ is considered movable if its preference list contains an alternative loop $L_k$ satisfying two conditions: (i) the spherical arc length $\Omega(L_k, d_i) \geq \Omega_{\text{floor}}$, and (ii) the increase in center distance $\mathrm{cd}(L_k, d_i) - \mathrm{cd}(L_j, d_i) \leq \Delta_{\text{max}}$. If any birth cannot be moved under these constraints, it becomes the keeper. If all births are movable, the one with the highest move cost (largest center distance increase to its best alternative) is designated as keeper.

Once the keeper is selected, each non-keeper birth is reassigned to its best available alternative that satisfies the $\Omega$ and center distance constraints and is not already assigned to another birth. The procedure is summarised in Algorithm 1. The code runs label collision up to 2 passes.

**Algorithm 1** Label Collision Resolution

**Require:** Initial assignment $\phi$, preference lists $\{\mathcal{P}_i\}$

**Ensure:** Refined assignment $\phi'$

```
$\phi' \leftarrow \phi$
for each label $L_j$ with $|B_j| > 1$ where $B_j = \{i : \phi'(i) = L_j\}$ do
  {Find keeper: birth that cannot be moved}
  keeper $\leftarrow \bot$
  for each $i \in B_j$ do
    canMove$_i$ $\leftarrow$ false
    for each $L_k \in \mathcal{P}_i[1:]$ do
      if $\Omega(L_k, d_i) \geq \Omega_{\text{floor}}$ and $\text{cd}(L_k, d_i) - \text{cd}(L_j, d_i) \leq \Delta_{\max}$ then
        canMove$_i$ $\leftarrow$ true
        break
      end if
    end for
    if $\neg$canMove$_i$ then
      keeper $\leftarrow i$
    end if
  end for
  if keeper $= \bot$ then
    keeper $\leftarrow \arg\max_{i \in B_j}$(move cost)
  end if
  {Move non-keepers to alternatives}
  for each $i \in B_j \setminus \{\text{keeper}\}$ do
    for each $L_k \in \mathcal{P}_i[1:]$ do
      if $L_k$ is free and $\Omega(L_k, d_i) \geq \Omega_{\text{floor}}$ and $\Delta\text{cd} \leq \Delta_{\max}$ then
        $\phi'(i) \leftarrow L_k$
        break
      end if
    end for
  end for
end for
return $\phi'$
```

**Same-Death Resolution.** When multiple births share the same death point and are assigned to the same loop, we keep the birth whose birth point is closest to the loop voxels and attempt to reassign the others to $\Omega$-qualified alternatives. Births without valid alternatives are marked as unpaired ($\phi(i) = \bot$).

**Strict Final Cleanup.** Any remaining collisions are resolved by selecting a single winner per label. For each collided label $L_j$ with births $B_j$, if birth $i$ has any $\Omega$-qualified candidate in its preference list, it is scored by center distance (lower is better); otherwise by spherical arc length (higher is better). The winner is the birth with the best score, and all others are marked unpaired.

##### 1.2.6 Parameters

| Parameter | Symbol | Value |
|---|---|---|
| k-NN neighbors | $k$ | 768 $\mu m$ |
| Minimum candidates | $\tau_{\min}$ | 20 $\mu m$ |
| $\Omega$ qualification floor | $\Omega_{\text{floor}}$ | 5.0 rad |
| Hard constraint tolerance | $\epsilon$ | 20 $\mu m$ |
| Max center-dist delta | $\Delta_{\max}$ | 18.0 $\mu m$ |
| Offset | offset | 25 $\mu m$ |

**Table 1.** Algorithm parameters for persistence pair matching.
